# Striatal D2-neurons gate D1-mediated associative encoding under uncertainty

**DOI:** 10.64898/2026.10.16.759515

**Authors:** Chelsea S Goulton, Christopher R Nolan, Nita R Roschanzamir, Melissa G Tran, Miriam Matamales, Jesus Bertran-Gonzalez

**Author notes:** These authors contributed equally: M. Matamales, J. Bertran-Gonzalez. **Contributions:** MM and JB-G conceived and de-signed the study. CSG, JB-G, and MM designed the experiments. CSG, JBG and NRR collected and analysed behavioural data. MM, CSG, MGT and NRR conducted histological procedures. CSG acquired and parametrised biosensor and optogenetic materials. MM performed confocal image acquisition and analysis. CSG collected fibre photometry data. CRN and CSG analysed fibre photometry data. CRN, CSG, MM and JBG developed data extraction and analysis code. CRN developed Behapy photometry analysis pipeline. MGT and CSG performed stereotaxic surgeries. JBG, CSG and CRN conducted statistical analyses. JBG, MM and CSG wrote the manuscript. **Competing interests:** The authors declare no competing interests.

## Abstract

Goal-directed actions are established through the encoding of predictive relationships between behaviour and its consequences, a process involving dopamine-dependent plasticity in the dorsal striatum. Early phases of learning entail complex associative landscapes from which useful action-outcome contingencies must be identified. Although dopamine processes are central to associative learning, the cellular mechanisms through which behaviourally relevant plasticity is selected from competing signalling events remain unknown. Here we show that D2-spiny projection neurons (D2-SPNs) are transiently recruited during the earliest stages of instrumental learning, when critical action-outcome contingencies emerge. We identify a dominant cell-to-cell transmodulation process by which D2-SPNs regulate dopamine-dependent signalling by constraining protein kinase A activity in D1-SPNs. This mechanism overrides extracellular dopamine signals and can be reproduced *in vivo* through calibrated optogenetic activation of cAMP signalling. Enhancing D2→D1-SPN transmodulation during uncertain reinforcement schedules durably improves action-outcome encoding, revealing a mechanism through which striatal networks promote relevant contingencies under associative ambiguity.

## Introduction

Associative learning enables organisms to identify useful predictive relationships between actions and their consequences, supporting adaptive behaviour. Yet the earliest stages of learning present a substantial challenge. Classical studies of instrumental learning highlighted how adventitious behaviours can be incidentally reinforced alongside causally effective actions^1–5^, exposing a noisy associative landscape from which functionally relevant contingencies must emerge. Contemporary theories describe acquisition as a process in which multiple candidate associations initially compete for behavioural control, while less informative relationships are progressively weakened or prevented^6–8^. Consequently, goal-directed learning is associated with increasing selectivity of instrumental performance. At the neural level, performance during acquisition is accompanied by a progressive reduction in activity pattern variability as neural responses stabilise around task-relevant representations^9–12^.

The dorsal striatum is central to instrumental learning and behavioural control^13^. Its principal neurons, dopamine D1- and D2-receptor-expressing spiny projection neurons (D1- and D2-SPNs), are densely intermingled within the striatum^14^, receive largely overlapping cortical inputs^15^, and contribute differently to adaptive behaviour^16–18^. The acquisition of goal-directed actions depends critically on intracellular signalling pathways that integrate dopamine and glutamate signals to drive long-term cellular adaptation^19–21^. Emerging evidence suggests that plasticity supporting the acquisition of goal-directed actions is prominent in D1-SPNs^22,23^. However, D1-SPN plasticity alone does not explain how behaviourally relevant contingencies are selected from the noisy associative landscape that is typical in early phases of learning.

Previous work demonstrated that D1- and D2-SPNs are linked through D2→D1-SPN transmodulation, a local mechanism through which activity in one population can influence plasticity-related processes in the other^24^. This organisation raises the possibility that D2-SPNs contribute to initial learning by gating which action-outcome contingencies are incorporated into D1-dependent plasticity.

Here we show that D2-SPNs are transiently recruited during the earliest stages of instrumental training and engage a dominant postsynaptic process that tightly regulates dopamine-dependent signalling in D1-SPNs. Using an optogenetic approach calibrated to emulate learning-related intracellular signalling dynamics, we demonstrate that selectively enhancing transmodulation at the time of reinforcement strengthens and stabilises action-outcome encoding. Together, these findings establish D2→D1-SPN transmodulation as a mechanism through which behaviourally relevant contingencies gain preferential access to D1-dependent plasticity during periods of associative uncertainty.

## Results

### Transitory engagement of D2-SPNs in early instrumental learning

Distinct territories of the dorsal striatum have been identified as key substrates for the associative plasticity underpinning instrumental learning. Dorsomedial striatal circuits preferentially support the acquisition of goaldirected action, whereas lateral regions become increasingly engaged as actions begin to automatise into habits^25–27^. In support, medio-lateral drifts of presynaptic corticostriatal and dopaminergic activity have been observed in the striatum as behavioural programs transition between associative structures^28–30^. To resolve the temporal and cellular organisation of learning-related territory shifts, we mapped transcriptionally activated spiny projection neurons (taSPNs) across multiple phases of instrumental training, distinguishing D1- and D2-SPN recruitment throughout the mediolateral extent of the striatum^24,31,32^ (Figure 1). Eight groups of D2-eGFP mice were exposed to different stages of instrumental conditioning in which lever pressing (action) produced food pellet delivery (outcome) under progressively increasing random ratio (RR) schedules of reinforcement. Instrumental performance improved steadily with training (Figure 1B), and responding became increasingly stereotyped from the RR10 stage onwards, consistent with the emergence of behavioural automatisation^31–34^ (Figure 1C). Maps of taD1-SPN distribution revealed strong transcriptional engagement of medial striatal areas during early training (1stC-RR5), which progressively shifted towards lateral territories in extended training groups (RR10-aRR20; Figure 1D, F). These learning-related changes were not observed in yoked control groups exposed to matched reward delivery in the absence of an action-outcome contingency. The progressive shift in D1-SPN recruitment from medial to lateral striatal territories coincided with the appearance of structured behavioural performance (cf. Figure 1C), revealing the temporal progression of learning-related territory reorganisation at the postsynaptic level.

**Figure 1.**
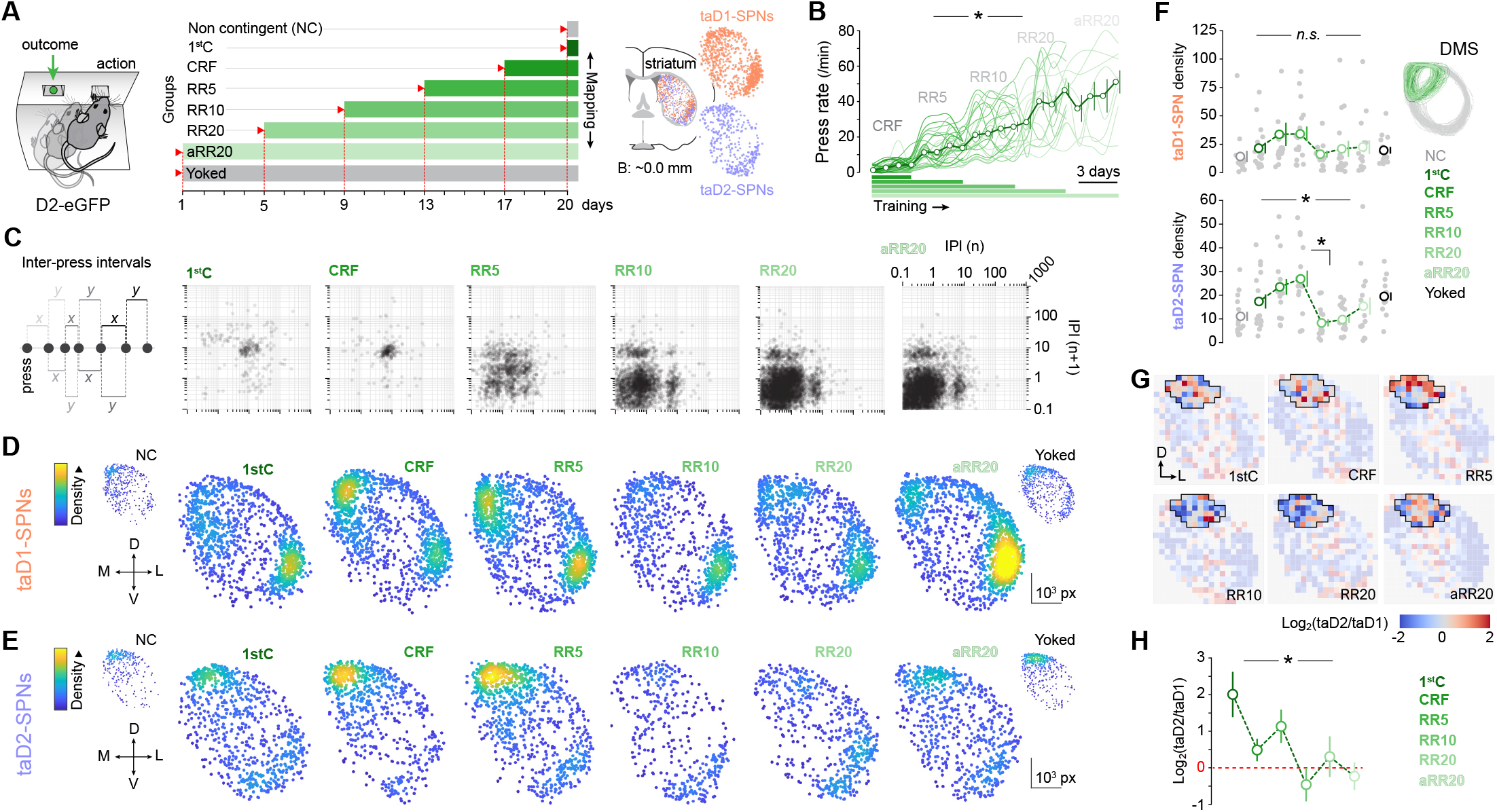
D2-SPN signalling is transiently engaged in medial striatal areas during early instrumental learning. **(A)** Eight groups of D2-eGFP were exposed to different levels of action (A; pressing a lever) → outcome (O; delivery of a food pellet) training (left), ranging from 1 to 20 days of training on an increasing random ratio schedule of reinforcement (centre). Transcriptionally active (ta) neurons displaying nucleosomal activation (phospho-histone-H3-immunoreactivity) were mapped and classified as D1- or D2-SPNs throughout the striatum (right). **(B)** Instrumental performance across training stages and groups. **(C)** Return maps of inter-press intervals (IPIs) displaying behavioural performance across phases of instrumental training. Left: each data point represents the time delay to its preceding (*x*; n) and succeeding (*y*; n + 1) behavioural element. **(D, E)** Digital reconstruction of taD1- (D) and taD2-SPNs (E) throughout the striatum of all animals in each training group (>13,500 taSPNs mapped). Scatterplots are stacked hemisections with pseudocoloured density highlighting clustering. **(F)** Quantification of taSPN density (taD1, top; taD2, bottom; nuclei/mm2 ) in dorsomedial striatal areas (DMS) delineated in each training group (left). **(G)** taD2/taD1-SPN ratio (log_2_) mesh analysis in whole striatum sections of each training group (20 × 20 grid, all mice). Higher values (red) indicate relative taD2-SPN enrichment, whereas lower values (blue) indicate taD1-SPN dominance. The area showing the greatest change is outlined in black.**(H)** Quantification of taD2/taD1-SPN ratios (log_2_) in high-change areas across training groups. Red dashed line (0) denotes equal proportion of taD2- and taD1-SPNs. Data in (B), (F) and (H) show mean values (dots) and SEM (error bars). ^*^, significant overall effect (black) and simple effect. *n*.*s*., not significant. See Table S1 for sample sizes and statistical analyses.

Unlike D1-SPNs, D2-SPNs did not exhibit a gradual medial-to-lateral redistribution. Instead, taD2-SPN recruitment was transient, with elevated densities in the dorsomedial striatum (DMS) during early training (1stC-RR5) that largely disappeared following extended training (RR10-aRR20; Figure 1E, F). We next mapped taD2/taD1-SPN ratios throughout the striatum to identify local territories in which activated D2-SPNs outnumbered activated D1-SPNs, a configuration predicted to favour D2→D1-SPN transmodulation^24,32^. This analysis revealed a prominent taD2-SPN dominance in dorsomedial territories that appeared during initial training stages (1stC-RR5; Figure 1G, H). These observations identify a selective enrichment of putative D2→D1-SPN transmodulatory events during the period in which new action-outcome contingencies are emerging.

### D2→D1-SPN transmodulation dominates molecular signalling in D1-SPNs

The emergence of D2-SPN-dominated territories during the earliest stages of instrumental learning was paradoxical, as this period is also associated with enhanced dopamine function and strong D1-SPN recruitment^22,35^. To investigate the functional significance of this organisation, we exploited the dual effects of pharmacological D2-receptor blockade at presynaptic and postsynaptic sites, which simultaneously causes elevation of striatal dopamine and recruitment of D2-SPN signalling^36,37^ (Figure S1). This allowed us to examine the consequences of D2→D1-SPN transmodulation under high levels of dopamine, approximating the neurochemical conditions that favour action-outcome encoding (Figure 2). In animals prepared for *in vivo* fibre photometry recordings of dopamine fluctuations (dLight1.1; Figure 2A), systemic administration of the D2-receptor antagonist raclopride (Rac) produced a prolonged elevation of extracellular dopamine in the DMS (Figure 2B). The same treatment induced robust intracellular signalling responses selectively in D2-SPNs, as indicated by strong nucleosomal activation (Figure 2C; Figure S2).

**Figure 2.**
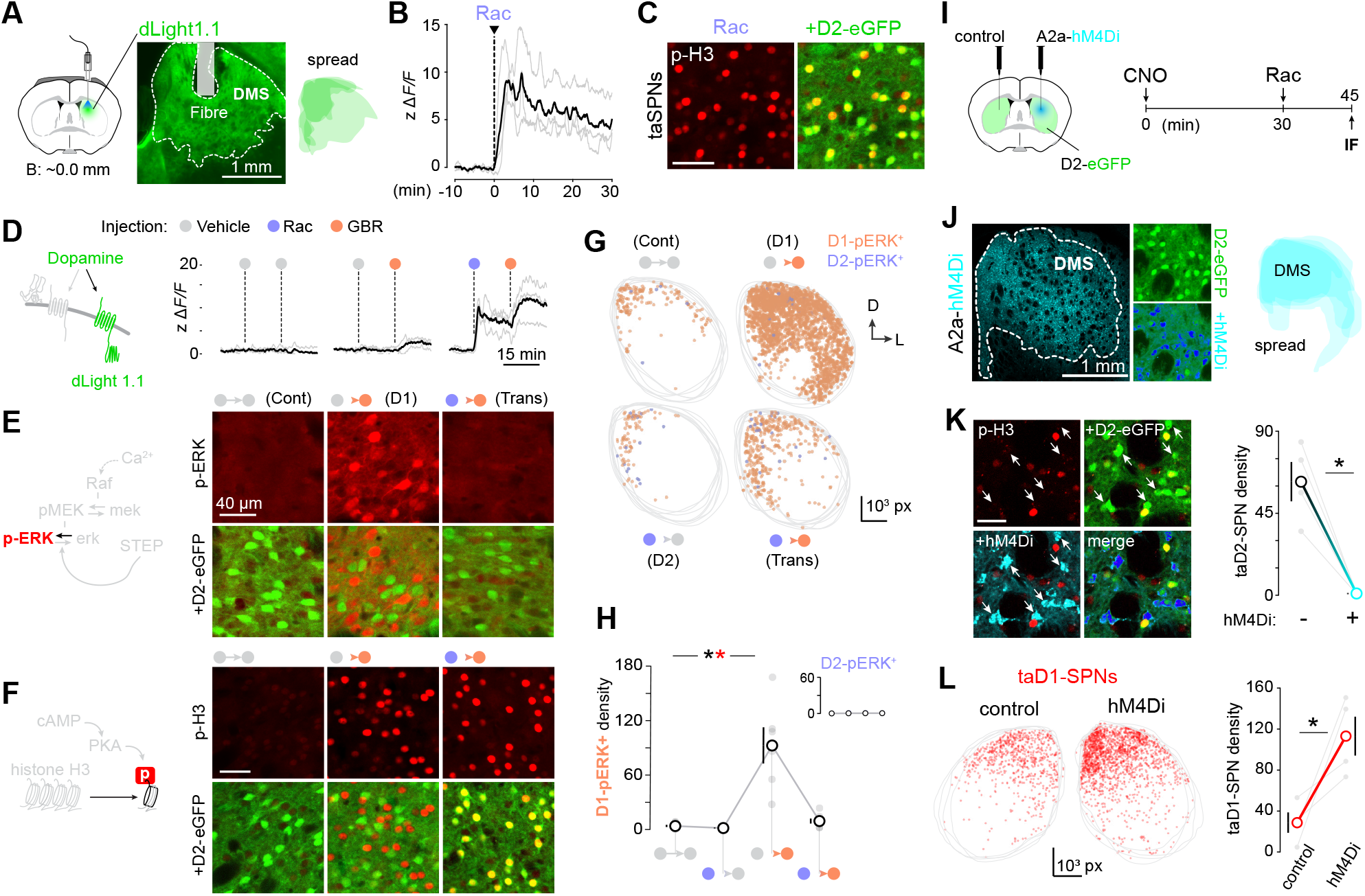
Pharmacologically-induced D2→D1-SPN transmodulation exposes a dominant postsynaptic process. **(A)** Mice were implanted with optic fibres into DMS for *in vivo* recording of extracellular dopamine (dLight1.1). **(B)** DMS dLight1.1 recordings before and after systemic administration of the D2-receptor antagonist Raclopride (Rac). **(C)** Confocal micrographs illustrating transcriptionally active SPNs displaying nucleosomal activation (immunoreactive for phosphorylated histone H3, p-H3) in the DMS of D2-eGFP mice collected 30 min after Rac administration. Co-localisation of p-H3 and eGFP identifies taD2-SPNs. **(D-F)** D2→D1-SPN transmodulatory effects were pharmacologically induced in the striatum of dLight1.1-bearing or D2-eGFP-expressing mice by combining a first injection of Rac (stimulates D2-SPN signalling) with a subsequent injection of GBR12783 (GBR, stimulates D1-SPN signalling) (Figure S1). Effects of various pharmacological conditions on dopamine release (D, dLight1.1 recordings), activation of cytonuclear signalling (E, p-ERK immunodetection) and nucleosomal activation (F, p-H3 immunodetection) in the DMS (Figure S2). **(G, H)** Mapping (G) and quantification (H) of p-ERK+ neurons classified as D1- and D2-SPNs across the striatal section in each pharmacological condition. **(I)** Left: unilateral expression of Cre-dependent hM4Di in the DMS of A2a-Cre/D2-eGFP mice. Right: pharmacogenetic approach to test cell-specific hM4Di-mediated inhibition on Rac-induced signalling responses in the striatum. **(J)** Selective expression of hM4Di in D2-SPNs of the DMS. **(K)** Effects of CNO administration on Rac-induced nucleosomal responses in hM4Di-expressing D2-SPNs (arrows). **(L)** Within-subjects comparison of taD1-SPNs in controlversus hM4Di-injected sides. Data in (H), (K) and (L) show mean values (dots) and SEM (error bars). ^*^, significant overall effect (black), simple effect (black) and interaction (red). See Table S1 for sample sizes and statistical analyses.

The dissociation between dopamine availability and D1-SPN signalling persisted when Rac was combined with GBR12783 (GBR), a dopamine transporter (DAT) blocker that further elevates extracellular dopamine levels (Figure 2D). Despite markedly elevated striatal dopamine, D1-SPNs remained largely silent, as indicated by the absence of p-ERK and p-H3, molecular markers typically observed in transcriptionally active D1-SPNs (Figure 2E-H, Figure S1, Figure S2). We next sought to dissociate the pre- and postsynaptic effects of Rac by chemogenetically preventing cAMP-dependent signalling in D2-SPNs within the DMS of A2a-Cre × D2-eGFP mice (Figure 2I). This manipulation was intended to leave Rac-induced dopamine elevations intact while selectively disrupting postsynaptic engagement of D2-SPNs, therefore bypassing D2→D1-SPN transmodulation. Confocal microscopy validated expression of the synthetic Gi-coupled GPCR hM4Di in D2-SPNs (Figure 2J) and confirmed that administration of CNO effectively prevented intracellular activity in hM4Di-expressing D2-SPNs (Figure 2K). Territories in which D2-SPN signalling was chemogenetically suppressed exhibited a marked increase in taD1-SPN recruitment relative to matched control hemispheres (Figure 2L). This indicated that preventing D2→D1-SPN transmodulation restored the ability of Rac-induced dopamine elevation to engage intracellular signalling in D1-SPNs. Together, these findings identify a dominant postsynaptic mechanism through which D2→D1-SPN transmodulation can override dopamine-dependent signalling in neighbouring D1-SPNs.

### D2→D1-SPN transmodulation spares Ca^2+^dynamics

Because the molecular markers examined above integrate both cAMP- and Ca^2+^-dependent signalling pathways, we next asked whether pharmacological D2→D1-SPN transmodulation influences intracellular Ca^2+^ levels, which are driven predominantly by excitatory corticostriatal and thalamostriatal glutamatergic transmission^38^ (Figure S1). Mice prepared for *invivo*fibre photometry recordings of Ca^2^d^+^ynamics in D1-SPNs (D1-GCaMP6m) were placed in an open field, where they received the pharmacological protocol used to engage D2→D1-SPN transmodulation (i.e., Rac-GBR), while locomotor activity was simultaneously monitored by video tracking (Figure S3A, B).

As expected, Rac induced a reduction in locomotion that did not prevent the hyperlocomotor effects of GBR (Figure S3C, D). D1-SPN intracellular Ca^2+^ fluctuations largely tracked behavioural activity across pharmacological conditions, decreasing following Rac administration and fluctuating following GBR injection (Figure S3E-H).

To investigate this relationship in greater detail, we quantified the temporal correspondence between behavioural pose and simultaneously recorded Ca^2+^ signals (Figure S4A, B). Behavioural variables consistently preceded optimally correlated Ca^2+^ signals by approximately 500 ms, regardless of the pharmacological state experienced by the animals (Figure S4C-G). Similar relationships between behavioural activity and Ca^2+^ dynamics were observed in D2-SPNs (Figure S5A-F). Together, these findings indicate that striatal Ca^2+^ dynamics remain tightly coupled to behavioural state across pharmacological conditions and are largely preserved despite pharmacological engagement of D2→D1-SPN transmodulation. Thus, the profound suppression of D1-SPN molecular signalling described upon D2-SPN activation could not be readily explained by changes in intracellular Ca^2+^ dynamics.

### D2-SPNs control cAMP-dependent signalling in D1-SPNs

The preservation of intracellular Ca^2+^ dynamics suggested that D2→D1-SPN transmodulation uncouples dopamine release from intracellular signalling through the cAMP-dependent arm of this pathway. To localise the signalling step at which this occurs, we used *in vivo* fibre photometry to monitor intracellular cAMP (dw-cADDis^39^) and PKA activity (ExRai-AKAR2^40^) in D1-SPNs of the DMS, together with extracellular dopamine measurements (dLight1.1^41^) (Figure 3A-C, Figure S6). When preceded by vehicle injection, dopamine transporter (DAT) blockade by systemic GBR triggered the canonical signalling responses expected in D1-SPNs (Figure S1). Elevated dopamine levels (Figure 3D, left) coincided with a burst of cAMP synthesis (Figure 3E, left) and a sustained increase in PKA activity that persisted for at least 30 min (Figure 3F, left). When GBR treatment was instead preceded by Rac administration, extracellular dopamine again increased in a stepwise manner, with each drug contributing a comparable elevation (Figure 3D, G). Remarkably, each dopamine elevation was accompanied by a corresponding pulse of intracellular cAMP, with cAMP levels rising and returning to baseline after each injection, consistent with tight regulation by the opposing activities of adenylyl cyclases and phosphodiesterases^42^ (Figure 3E, H). Despite this preserved coupling between dopamine and cAMP, prior Rac administration strongly attenuated the increase in PKA activity normally produced by GBR (Figure 3F, I). Thus, D2→D1-SPN transmodulation left dopamine release and cAMP production largely intact while disrupting the propagation of signalling to downstream PKA-dependent processes.

**Figure 3.**
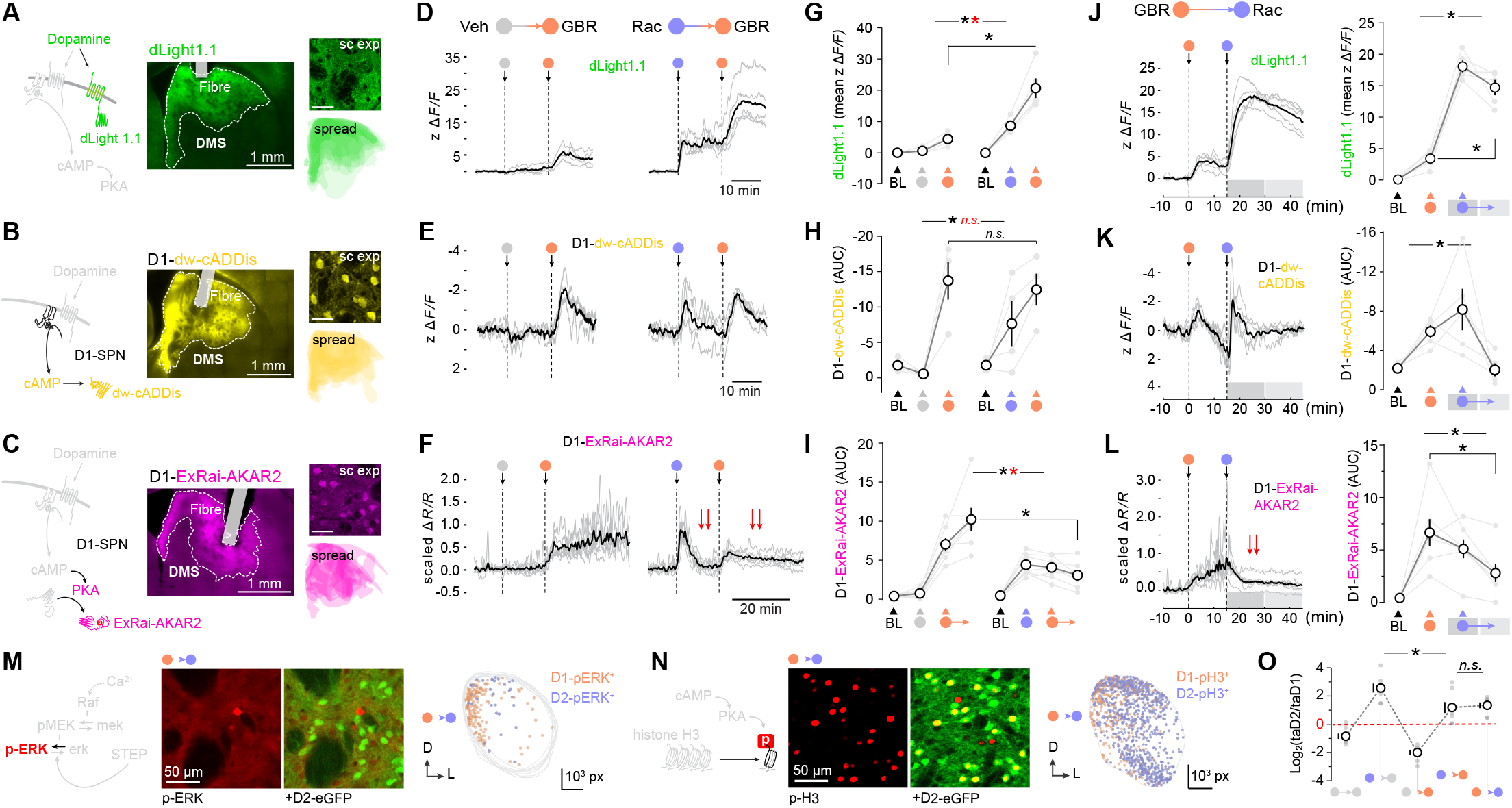
D2→D1-SPN transmodulation intercepts ongoing cAMP/PKA signalling in D1-SPNs. **(A-C)** Mice were implanted with optic fibres into the DMS for *in vivo* recording of (A) extracellular dopamine (dLight1.1 in WT mice; green), (B) intracellular cyclic AMP (cAMP; DIO-dw-cADDIs in D1-Cre mice; yellow) and (C) protein kinase A activity (PKA; Flex-ExRai-AKAR2 in D1-Cre mice; magenta). For each biosensor, insets show subcellular expression (sc exp, top) and expression spread (bottom) observed in the DMS. Inset scale bars: 40 μm. **(D-I)** Fibre photometry recordings (D-F) and quantification (G-I) of striatal dopamine (G, G), D1-cAMP (E, H) and D1-PKA activity (F,I) signals in mice exposed to pharmacologically-induced D2→D1-SPN transmodulation (Rac→GBR). **(J-L)** *In vivo* recording (left) and quantification (right) of dopamine (J), cAMP (K) and PKA activity (L) following reversal of pharmacological treatment order (GBR→Rac). **(M, N)** Confocal micrographs showing the effects of GBR→Rac treatment on downstream cytonuclear (p-ERK; M) and nucleosomal (p-H3; N) signalling in D1- and D2-SPNs. **(O)** Quantification of taD2/taD1-SPN ratios (log_2_) throughout the striatum across treatment groups. Red dashed line (0) denotes equal proportions of taD2- and taD1-SPNs. Double red arrows in (F) and (L) indicate abrupt reductions in PKA-reporter activity. Data show individual values (light grey), mean values (dots) and SEM (error bars). ^*^, significant overall/simple effect (black) and interaction (red). *n*.*s*., not significant. See Table S1 for sample sizes and statistical analyses.

Further evidence localising this effect to the level of PKA emerged from responses to Rac alone. Although D2-receptor blockade caused parallel elevations of dopamine, cAMP and PKA activity, the latter rapidly collapsed back to baseline approximately 10 min later (Figure 3F; Figure S7). This unexpected dissociation suggested that, beyond preventing it, D2→D1-SPN transmodulation can also interrupt ongoing PKA signalling. To test this possibility directly, we reversed the order of drug administration, allowing downstream signalling to be established before Rac administration (Figure 3J-N). A first injection of GBR triggered canonical dopamine/cAMP/PKA signalling, whereas a subsequent injection of Rac further elevated extracellular dopamine (Figure 3J) and elicited a second pulse of cAMP synthesis (Figure 3K), yet strongly and durably suppressed established PKA activity (Figure 3L). D2-SPN activation also downregulated ongoing PKA-mediated phosphorylation of target proteins in D1-SPNs, as reflected by reduced cyto-nuclear p-ERK (Figure 3M) and nucleosomal p-H3 levels (Figure 3N). Consistent with these observations, quantification of taD2/taD1-SPN ratios revealed a robust taD2-SPN dominance throughout the striatum whenever Rac was present, irrespective of the pharmacological protocol employed (Figure 3O). Together, these findings demonstrate that D2-SPN-mediated transmodulation can rapidly intercept already activated PKA-dependent signalling in D1-SPNs despite preserved upstream dopamine and cAMP dynamics (Figure S1).

### Real-time manipulation of D2→D1-SPN transmodulation

So far, D2→D1-SPN transmodulation had been examined using pharmacological manipulations that simultaneously increased dopamine availability and D2-SPN engagement. We next sought to isolate the postsynaptic component of this process by selectively inducing cAMP signalling in D2-SPNs. To achieve this, we used biPAC, a blue-light-sensitive bacterial adenylyl cyclase optimised for minimal activity in the dark^43–45^. In A2a-Cre × D2-eGFP mice bearing bilateral optical implants, Cre-dependent biPAC was expressed unilaterally in D2-SPNs of the DMS, while the contralateral hemisphere received Cre-dependent mCherry (Figure 4A). Unlike mCherry control hemispheres, intermittent blue-light stimulation resulted in p-H3 induction selectively in D2-SPNs within biPAC-expressing territories (Figure 4B). Thus, under these stimulation conditions, biPAC successfully engaged the full cAMP/PKA signalling column in D2-SPNs, from cytoplasmic activation to nuclear transcriptional responses (Figure S1).

**Figure 4.**
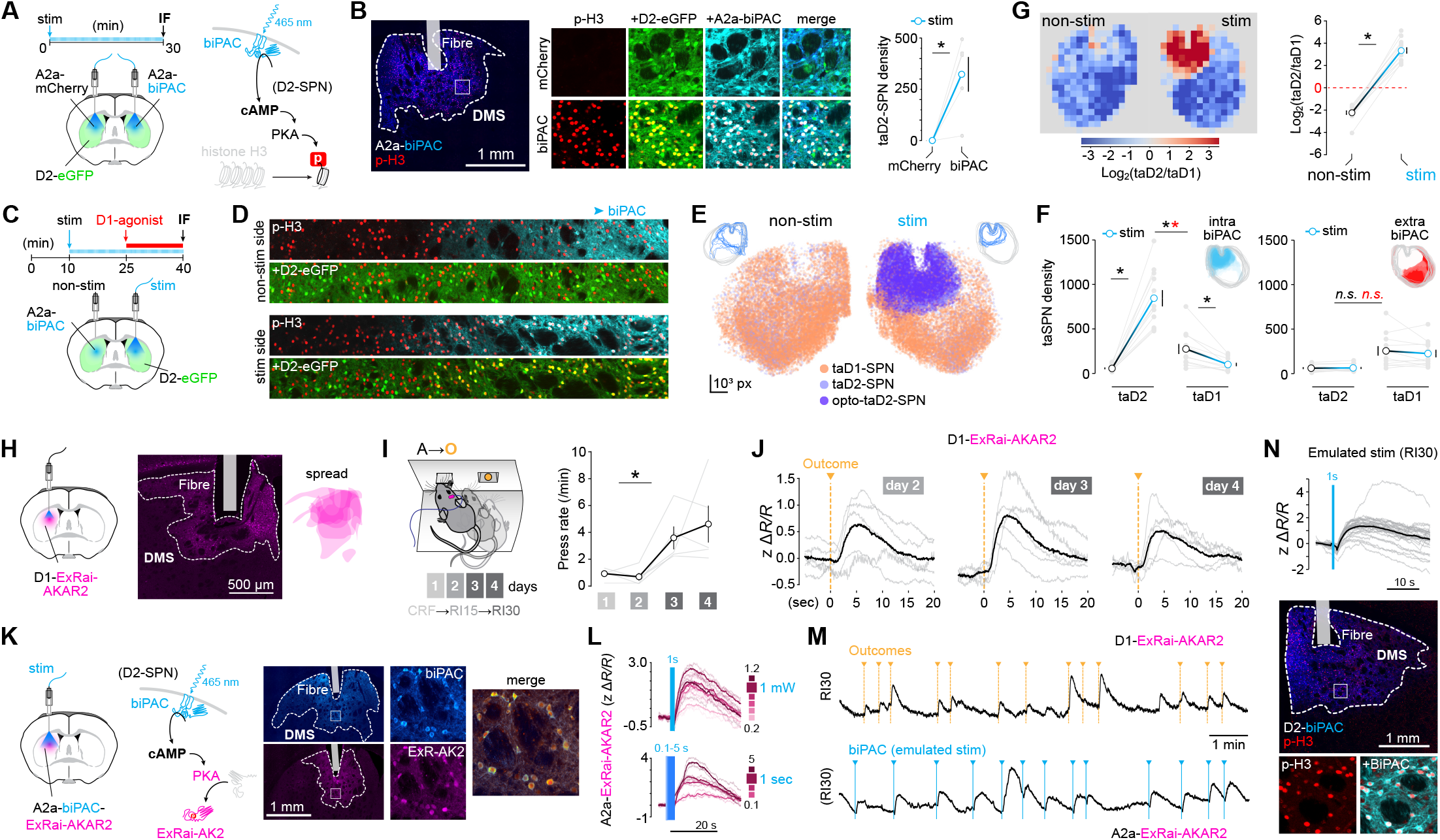
Optogenetic control of D2→D1-SPN transmodulation. **(A)** A2a-Cre/D2-eGFP mice received unilateral injections of Cre-inducible biPAC (allows optostimulation of cAMP signalling, right) and contralateral injections of Cre-inducible mCherry (control) in the DMS, followed by bilateral optic fibre implantation. Mice received repeated optical stimulation (1 mW, 1 s on, 10 s off) for 30 min and were subsequently perfused for immunofluorescence analyses. **(B)** Confocal micrographs and quantification of transcriptionally active SPNs displaying nucleosomal activation (p-H3) in mCherry- and biPAC-expressing DMS areas. Images show co-localisation of p-H3+ neurons with eGFP (identifying D2-SPNs) and AAV-mediated fluorophore (mCherry, top; biPAC-mKate2, bottom). Right, density of co-localised p-H3+ nuclei (taD2-SPNs) in paired mCherry- and biPAC-expressing hemispheres. **(C)** A2a-Cre/D2-eGFP mice bearing bilateral Cre-dependent biPAC and optic fibre implants in the DMS received unilateral optical stimulation (as in (A)) and systemic D1-receptor agonist injections before tissue collection for immunofluorescence. Stimulation: t = 10-40 min; injection: 1.5 mg kg^−1^ chloro-APB at t = 25 min. **(D)** Wide-field confocal micrographs showing taSPNs across biPAC-negative to biPAC-positive transition zones in stimulated (bottom) and non-stimulated (top) hemistriata of the same animal (Figure S8). **(E)** Digital reconstruction of taD1- and taD2-SPNs throughout non-stimulated and stimulated hemistriata following the protocol shown in (C). taD2-SPNs were further classified into optically stimulated taD2-SPNs (opto-taD2-SPNs) based on detectable biPAC expression. Insets show stacked hemistriata and contours of biPAC-expressing territories. Each side shows three consecutive hemisections from each of four mice, stacked for visualisation (>40,000 taSPNs mapped). **(F)** Quantification of taSPN density across cell types and non-stimulated (black) versus stimulated (blue) hemistriata in biPAC-expressing (intra) and non-expressing (extra) areas. **(G)** taD2/taD1-SPN ratio (log_2_) mesh analysis in non-stimulated and stimulated hemistriata (20 × 20 grid, one animal). Right, quantification of taD2/taD1-SPN ratios (log_2_) in biPAC-expressing regions of non-stimulated and stimulated sides across all mice. **(H)** D1-Cre mice injected with Cre-inducible ExRai-AKAR2 were unilaterally implanted with optic fibres into the DMS for *in vivo* recording of PKA activity in D1-SPNs. **(I)** Implanted mice underwent rapid A→O learning under an increasing random interval (RI) schedule of reinforcement over 4 days. **(J)** Individual (light grey) and average (black) traces of PKA-dependent activity aligned to delivery of instrumentally earned outcomes across RI training. **(K)** Co-expression of Cre-inducible biPAC and ExRai-AKAR2 in DMS D2-SPNs of A2a-Cre mice enabled simultaneous optical stimulation of cAMP and recording of PKA activity *in vivo* (right). **(L)** biPAC stimulation conditions were calibrated to mimic behaviourally observed PKA responses in D2-SPNs by testing a range of stimulation powers (top) and durations (bottom). **(M)** Example comparison of naturally occurring (top, D1-SPNs) and biPAC-evoked (bottom, D2-SPNs) PKA signalling dynamics recorded during A→O learning and RI30-emulated stimulation, respectively. **(N)** biPAC-evoked PKA activity (ExRai-AKAR2; top) and nucleosomal activation (p-H3; bottom) in D2-SPNs of A2a-Cre mice expressing Cre-dependent biPAC in the DMS following emulated RI30 stimulation (1 mW, 1 s pulses). Data in (F), (G) and (I) show individual values (light grey), mean values (dots) and SEM (error bars). ^*^, significant overall/simple effect (black) and interaction (red). *n*.*s*., not significant. See Table S1 for sample sizes and statistical analyses.

To test transmodulation under conditions restricted entirely to postsynaptic signalling, D2-SPN cAMP activity was triggered optogenetically through biPAC, whereas D1-SPN signalling was induced pharmacologically using the D1-receptor agonist chloro-APB^24^ (Figure 4C). This approach removed presynaptic dopamine regulations from the experimental design while maintaining controlled activation of both signalling systems. In non-stimulated biPAC-expressing hemistriata, D1-receptor agonist induced robust nucleosomal responses predominantly in eGFP-negative neurons, corresponding to D1-SPNs^46,47^ (Figure 4D, E; Figure S8). This distribution was maintained across both extra- and intra-biPAC territories. By contrast, optical stimulation produced a striking redistribution of nucleosomal activation. Whereas p-H3 induction remained largely restricted to D1-SPNs in extra-biPAC territories, stimulation of biPAC-expressing regions reduced D1-SPN activation and promoted robust p-H3 induction in D2-SPNs (Figure 4D-F; Figure S8). Consistent with this shift in activation, taD2/taD1-SPN mapping revealed that biPAC stimulation locally transformed D1-SPN-biased territories into D2-SPN-enriched regions (Figure 4G). These findings establish that D2→D1-SPN transmodulation can be induced through strictly postsynaptic manipulations and demonstrate that selective activation of cAMP signalling in D2-SPNs is sufficient to initiate transmodulatory effects in neighbouring D1-SPNs.

Having validated biPAC as a selective approach for engaging D2→D1-SPN transmodulation *in vivo*, we next calibrated optogenetic induction of D2-SPN cAMP signalling against endogenous signalling signatures observed during instrumental learning. D1-Cre mice expressing ExRai-AKAR2 in D1-SPNs of the DMS were trained on a short random-interval (RI) procedure, allowing signalling dynamics to be examined during the earliest stages of contingency encoding (Figure 4H, I). Across training, reward delivery consistently induced transient increases in PKA activity that peaked ∼5 s after outcome delivery and returned to baseline within approximately 20 s (Figure 4J). To reproduce these dynamics in D2-SPNs, we co-expressed biPAC and ExRai-AKAR2 in the DMS of A2a-Cre mice, enabling simultaneous optogenetic control of cAMP and real-time measurement of PKA activity (Figure 4K). Following systematic testing of stimulation parameters, 1-s pulses delivered at 1 mW most closely reproduced naturally occurring reward-evoked PKA responses (Figure 4L). These conditions enabled optogenetic emulation of prototypical RI30-associated cAMP/PKA signalling dynamics in D2-SPNs (Figure 4M). RI30-emulated stimulation induced sparse nucleosomal activation in D2-SPNs (Figure 4N), providing a physiologically grounded approach for generating discrete D2→D1-SPN transmodulation events during instrumental learning.

### Upregulation of D2→D1-SPN transmodulation strengthens associative encoding during instrumental learning

RI-driven stimulation was next used to increase D2→D1-SPN transmodulation during instrumental learning and assess its impact on associative encoding (Figure 5). A2a-Cre mice bilaterally expressing biPAC in DMS D2-SPNs received outcome-triggered optical stimulation, such that each instrumentally earned reward initiated a D2→D1-SPN transmodulation event (Figure 5A; cf. Figure 4M, N). Control mice expressed mCherry and received identical training and stimulation protocol. Viral expression was largely confined to medial striatal territories in both groups (Figure 5B). Mice were trained on a first action-outcome contingency (A1→O1) using increasing random-interval (RI) schedules, a procedure that prolongs the associative uncertainty characteristic of early instrumental learning^48,49^ (Figure 5C, left). Both groups acquired the instrumental response and completed all training sessions, but biPAC-expressing mice displayed lower overall lever-press rates than controls, reaching asymptotic performance by the fourth training day (Figure 5C, right). Subsequent locomotor testing demonstrated that these differences could not be explained by motor suppression or catalepsy-like effects (Figure S9), which are typically associated with prolonged stimulation of D2-SPNs^50^. Closer examination of instrumental response structure revealed that biPAC mice maintained highly focused patterns of task-relevant lever pressing throughout training (Figure 5D), suggesting enhanced allocation of behaviour towards actions most closely linked to reinforcement. To directly evaluate associative encoding of A1→O1, we assessed sensitivity to outcome devaluation^13,51^. Mice were tested under extinction following specific satiety on either O1 itself (O1 devalued) or regular chow (O1 valued) (Figure 5E, left). Although both groups pressed more when O1 remained valued (Figure 5E, right), biPAC mice displayed substantially greater sensitivity to devaluation, largely because they more effectively suppressed responding when the outcome’s value was reduced (Figure 5F). These findings indicate that optogenetic amplification of D2→D1-SPN transmodulation strengthens encoding of the instrumental contingency acquired during training.

**Figure 5.**
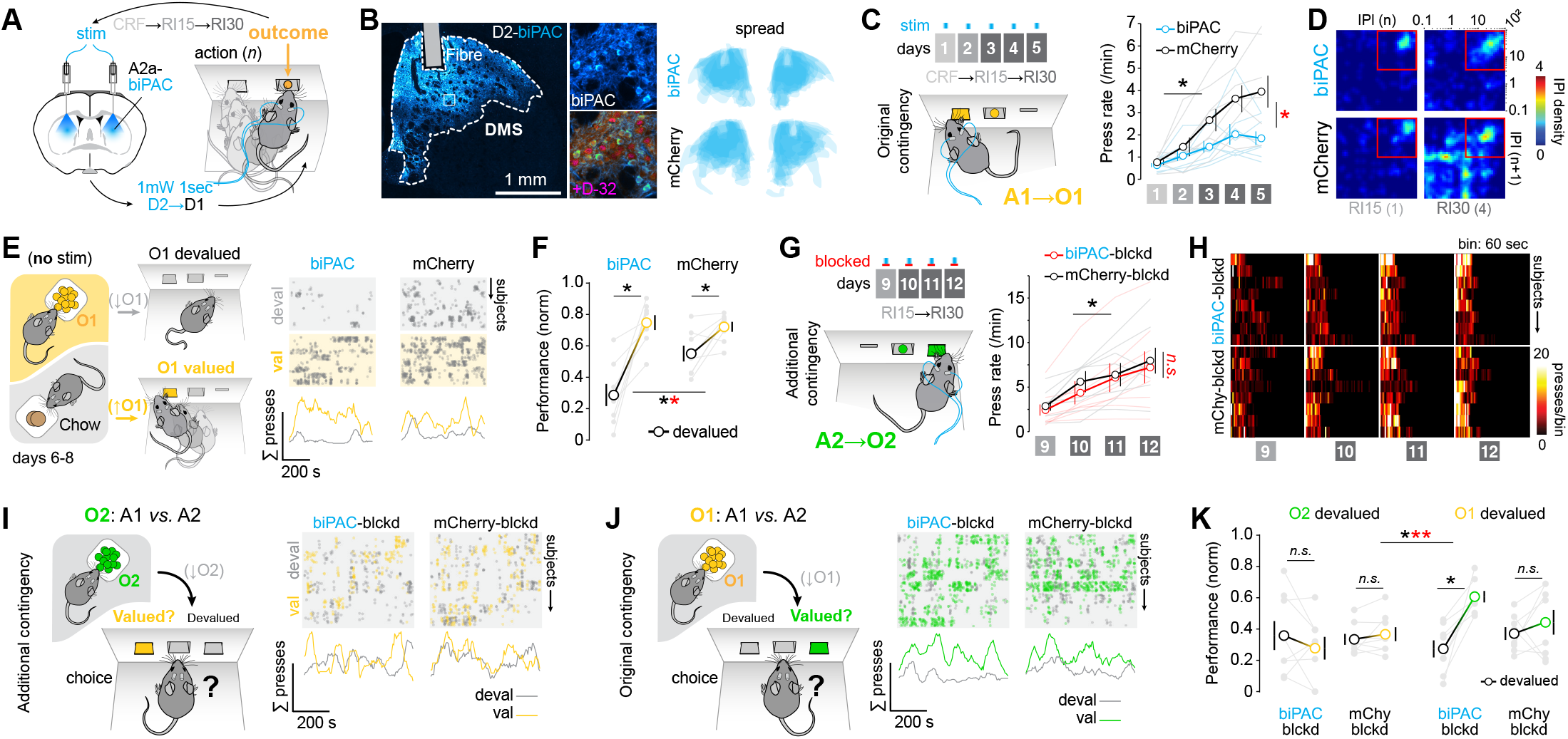
Reinforcement-coupled D2→D1-SPN transmodulation strengthens action→outcome encoding. **(A)** A2a-Cre mice bilaterally expressing Cre-dependent biPAC or mCherry and bearing bilateral optic fibre implants underwent looped optogenetic stimulation linking instrumentally earned outcomes to discrete D2→D1-SPN transmodulation events. **(B)** AAV expression in SPNs was confirmed by DARPP-32 (D-32) immunostaining. Right, extent of viral expression within bilateral DMS territories across all mice. **(C)** biPAC- and mCherry-expressing mice were trained on an initial action→outcome contingency (A1→O1; original contingency) under a five-phase random-interval (RI) schedule, during which delivery of earned outcomes triggered optical stimulation. Right, lever press rates across A1→O1 training. **(D)** Return maps of inter-press intervals (IPIs) showing behavioural performance during early and late RI training phases in each group. Red squares denote regions associated with efficient instrumental responding. **(E)** The strength of the A1→O1 contingency acquired during training was assessed using an outcome devaluation procedure in which instrumental performance was tested following satiation on either O1 or chow in a separate chamber. Right, distribution of lever presses (grey dots) in individual mice following satiation on O1 (devalued, grey trace) or chow (valued, yellow trace). **(F)** Quantification of instrumental performance during devalued (black) and valued (yellow) tests. **(G)** A second, distinct action→outcome contingency (A2→O2) was trained over a further four days of RI reinforcement. Optical stimulation was blocked during this phase in both groups (biPAC-blocked and mCherry-blocked). Right, lever press rates across A2→O2 training. **(H)** Lever-press density across sessions of A2→O2 training. **(I, J)** The strength of the recent (I, A2→O2) and original (J, A1→O1) contingencies was assessed using outcome-specific devaluation. Mice were sated on either O2 or O1 outcome before a two-lever choice test. Right panels show the distribution of lever presses on devalued (grey) and valued (coloured) levers in each group. **(K)** Quantification of choice performance across devaluation tests (A2→O2 versus A1→O1) and groups (biPAC-blocked versus mCherry-blocked). Data in (C), (F), (G) and (K) show individual values (light grey), mean values (dots) and SEM (error bars). Performance in (F) and (K) is lever-press rate normalised to the final day of training. Raster plots in (E), (I) and (J) show data from all mice organised from top to bottom. ^*^, significant overall/simple effect (black) and interaction (red). *n*.*s*., not significant. See Table S1 for sample sizes and statistical analyses.

We next examined whether this enhancement persisted as animals accumulated further associative learning. Following completion of the first devaluation test, mice were trained on an additional action-outcome contingency (A2→O2) under equivalent RI schedules, but with optical stimulation disabled in both groups (blocked) (Figure 5G, left). Under these conditions, all mice rapidly acquired the new response and displayed indistinguishable training performance (Figure 5G, right) and behaviour structuring (Figure 5H). We then compared the strength of the original and newly acquired contingencies using outcome-specific devaluation followed by choice tests, in which both levers were available simultaneously but no outcome was delivered^52,53^. We focused our analysis on the choice performance (A1 vs. A2) displayed immediately after devaluation of O2 (tests integrity of A2→O2; Figure 5I) or O1 (tests integrity of A1→O1; Figure 5J). We found that neither group displayed a clear choice upon devaluation of O2 (Figure 5I, K), suggesting that training of the additional contingency was either insufficient or was overshadowed by the previously acquired contingency^52,53^. By contrast, biPAC mice strongly biased performance following O1 devaluation, whereas control mice did not (Figure 5J, K). This finding indicates that the original A1→O1 contingency remained highly discriminable and accessible despite acquisition of an additional contingency and several days of intervening training. Together, these findings indicate that D2→D1-SPN transmodulation during acquisition promoted the selective incorporation and long-term stabilisation of action-outcome contingencies, rendering them resistant to disruption by subsequent learning.

## Discussion

In this study we identify a previously unrecognised role for D2→D1-SPN transmodulation in contingency encoding. We show that D2-SPNs are transiently recruited within dorsomedial striatal territories during early instrumental learning, engage a dominant postsynaptic process that hijacks dopamine-dependent signalling in D1-SPNs, and can be harnessed optogenetically to strengthen and stabilise action-outcome encoding in noisy associative environments. Together, these findings identify D2→D1-SPN transmodulation as a mechanism through which behaviourally relevant contingencies gain preferential access to D1-dependent plasticity during periods of associative uncertainty.

One intriguing feature of D2→D1-SPN transmodulation is its dominance over striatal dopamine dynamics. Rather than simply reflecting dopamine availability^46,54^, intracellular signalling in D1-SPNs can be overridden by neighbouring D2-SPNs. This counterintuitive arrangement has important implications for contemporary models of reward prediction error encoding because it positions D2-SPNs as local arbiters of dopamine-dependent plasticity. Within the cytoarchitecturally homogenous environment of the striatum, such a mechanism may be particularly advantageous for amplifying rapid changes in dopaminergic activity sweeping across the striatal continuum during periods of intensive associative encoding^30^. Despite evidence that negative prediction errors associated with inhibitory learning are conveyed by transient pauses in dopaminergic activity^55,56^, the cellular mechanisms through which such brief reductions in dopamine availability are translated into enduring modifications of striatal plasticity remain poorly understood. We report that D2-SPNs, which are sensitive to dopamine reductions, can rapidly intercept ongoing dopamine-dependent signalling in neighbouring D1-SPNs. This mechanism provides a means through which local striatal circuits may register and amplify transient decreases in dopamine signalling, thereby regulating the incorporation of new information into ongoing learning. Our finding that transmodulation-assisted contingencies were acquired more robustly and remained resilient to subsequent interference arising from additional learning raises the possibility that D2→D1-SPN transmodulation contributes to the progressive elimination of competing associative elements, effectively narrowing the pool of candidates available for reinforcement. In complex associative settings such as early instrumental learning, D2→D1-SPN transmodulation may therefore serve an associative pruning function that determines which dopamine-dependent plasticity events are permitted to proceed.

The role of D2-SPNs in associative learning and goal-directed action has become increasingly recognised^23,24,32,57^, yet the specific cellular activities that expose learning-related plasticity processes require careful consideration. Our results show that Ca^2+^dynamics, widely used as a proxy of neuronal activity, do not capture D2→D1-SPN transmodulation and instead closely track ongoing behaviour, likely largely reflecting corticostriatal excitatory drive associated with motor execution^58^. By contrast, intracellular signalling pathways linked to dopamine modulation, particularly the cAMP/PKA cascade^19,46,54,59,60^, provided robust signatures of transmodulation throughout the striatum. These pathways sit at major points of convergence for diverse neuromodulatory signals and, by regulating nucleosomal responses and gene expression, are likely to provide a more faithful readout of associative learning than measures of neuronal activity alone^61^.

Our findings further reveal a striking form of horizontal communication between neighbouring SPNs. Selective induction of cAMP signalling in D2-SPNs by targeted optogenetics was sufficient to suppress D1R/cAMP/PKA-dependent signalling in nearby D1-SPNs. This observation suggests that transmodulation operates by altering the capacity of neighbouring neurons to undergo plasticity, shifting the balance between kinase- and phosphatase-dependent processes despite preserved upstream dopamine signalling. While the molecular identity of the transmodulatory signal remains unknown, the observation that molecular inhibition of D1-SPNs extends beyond areas of biPAC-photoactivated D2-SPNs (Figure S8) suggests that locally sourced diffusible signals may convey these effects. Given the considerable neurochemical complexity of the striatum^62^, particularly during the encoding of goal-directed learning^18,63^, multiple candidate signals are conceivable and future work will be required to identify them. Collectively, these results provide a new perspective on striatal learning processes by showing that postsynaptic transmodulatory interactions between D1- and D2-SPNs can constrain the learning consequences of elevated extracellular dopamine, thereby biasing reinforcement towards causally effective actions. More broadly, these findings call for a revised view of striatal function in which D1- and D2-SPNs operate as integrated partners rather than as purely opposing pathways. In this framework, intracellular signalling processes within local striatal circuits become critical determinants of how dopamine dynamics are translated into learning.

### Resources

Source data underlying all Figures and Supplemental Figures have been deposited in Figshare (DOI: https://doi.org/10.6084/m9.figshare.34465431).

Raw data for reanalysis and reuse have been deposited in Figshare and will be made publicly available upon publication (DOI: https://doi.org/10.6084/m9.figshare.33969307).

All custom code used for data extraction, processing, analysis and visualisation in this study has been deposited in GitHub and will be made publicly available upon publication (https://github.com/NML-NeuroModuLab/umbrella).

The fibre-photometry analysis process using behapy is publicly available via GitHub (https://github.com/crnolan/behapy).

All code and data are available from the lead contact on request.

Supplementary material is attached to this preprint (https://www.biorxiv.org/content/10.64898/2026.10.16.759515v2.supplementary-material).

## Methods

### Experimental model and study participant details Animals

All experimental procedures were approved by the Animal Care and Ethics Committee of the University of New South Wales and conducted in accordance with the Australian Code for the Care and Use of Animals for Scientific Purposes (2013; updated 2021). Adult mice (>3 months old) of both sexes were used throughout the study. Experiments used C57BL/6J mice obtained from Australian BioResources (Moss Vale, Australia) or transgenic mice maintained on a C57BL/6J-Ausb background. The transgenic lines, originally obtained from the Mutant Mouse Resource and Research Centers, were D2-eGFP, which expresses eGFP in D2-SPNs; A2a-Cre, which expresses Cre recombinase in D2-SPNs; and D1-Cre, which expresses Cre recombinase in D1-SPNs. Experimental animals included D2-eGFP F1 hybrids, A2a-Cre::D2-eGFP double-transgenic mice, and inbred D1-Cre and A2a-Cre mice. Mice were housed in individually ventilated cages within a climate-controlled colony room maintained on a 12-h light/dark cycle. Where possible, littermates from the same home cage were allocated across different experimental groups.

### Method details Surgery

#### General surgical procedures

Mice were anaesthetised in an induction chamber with 4% isoflurane (Laser Animal Health, Pharmachem, Australia) delivered in oxygen (0.5 L/min) and placed in a stereotaxic frame (Kopf Instruments) fitted with a nose cone supplying a continuous flow of oxygen/isoflurane mixture (1 L/min). The depth of anaesthesia was monitored by assessing pedal reflexes. A pre-operative subcutaneous injection of meloxicam (5 mg/kg) was administered to provide peri-operative analgesia. At the end of the procedure, mice were placed individually in recovery cages containing a warming pad until they regained normal mobility and were able to maintain body temperature. Experimental procedures commenced following a minimum 3-week recovery period.

#### rAAV injections

All rAAV-injections were targeted to the dorsomedial striatum (DMS; AP+0.13mm, ML 1.75 relative to bregma; DV 2.6mm below skull surface). For stereotaxic delivery, a midline scalp incision (∼1.5 cm) was made to expose the skull and a small craniotomy was made at the specified coordinates. Glass capillaries (3.5”, Drummond Scientific #3-000-203-G/X) were pulled using a micropipette puller (Sutter Instrument), back-filled with mineral oil and mounted in a nanoliter microinjector (RWD). To minimise viral deposition during pipette insertion, the injector was retracted by 20 nL to create negative pressure and the pipette tip was briefly inserted into an agar gel block prior to use. The micropipette was then slowly lowered vertically through the craniotomy to the target depth. Following a 2 min settling period, virus was injected at a rate of 1 nL/s. Five minutes after completion of the injection, the pipette was slowly withdrawn. After all injections were completed, the incision was closed with surgical silk suture and sealed with tissue adhesive (Vetbond, 3M).

#### Optical implants

Following skull preparation and viral injection as described above, additional holes were drilled for anchoring screws placed anterior and posterior to the target sites, with two screws used for unilateral implants and three screws used for bilateral implants. Optical fibres were lowered gradually (200 μm every 15 s) and positioned either 100 μm above the injection site for fibre photometry experiments or 200 μm above the injection site for optogenetic experiments. Fibres were secured using cyanoacrylate adhesive (Loctite 454, Loctite), which was rapidly cured with a small amount of accelerator (Zip Kicker, Zap) and reinforced with a cap of two-part acrylic dental cement (Castapress, Vertex Dental) to stabilise the implant and seal the surgical site. For fibre photometry experiments, optical-fibre implants with a 400 μm core, 0.66 numerical aperture, 3.5 mm fibre length, and M2 screw connector were used (Doric Lenses). For optogenetic experiments, optical-fibre implants with a 400 μm core, 0.50 numerical aperture, 1.25 mm bare ceramic ferrule, and custom 3.5 mm fibre length were used (RWD).

### Viral vectors and titres

Recombinant adeno-associated viral (rAAV) vectors were used throughout this study. The following vectors were obtained as packaged viral preparations from Addgene: AAV1.Syn.Flex.GCaMP6m.WPRE.SV40^67^ (7.7 × 10^12^ vg mL^−1^), AAV5-CAG-dLight1.1^41^ (9 × 10^12^ vg mL^−1^), AAV5-hSyn-DIO-hM4D(Gi)-mCherry (2.4 × 10^13^ vg mL^−1^), and AAV5-hSyn-mCherry (1.4 × 10^13^ vg mL^−1^). The remaining vectors were generated from Cre-dependent AAV transfer plasmids and custom packaged as indicated. For cADDis, pAAVhSyn-DIO-cADDis-DW-green.WPRE (Montana Molecular) was packaged as AAV1-hSyn-DIO-cADDis-DW-green.WPRE (1.25 × 10^13^ vg mL^−1^; VGEF). For ExRai-AKAR2^68^, the sensor sequence was obtained from pcDNA3.1(+)-ExRaiAKAR2 (Addgene), the Cre-dependent transfer plasmid pAAV-FLEX-syn1-ExRaiAKAR2-WPRE was synthesised by VectorBuilder, and the resulting vector was packaged as AAV2/1-FLEX-syn1-ExRaiAKAR2-WPRE (1.25 × 10^13^ vg mL^−1^; VVC). For biPAC^44^, pAAV-DIO-EF1a-mKate2-biPAC (Addgene) was packaged as AAV5-DIO-EF1a-mKate2-biPAC (1.4 × 10^13^ vg mL^−1^; VVC).

### Pharmacological manipulations

Raclopride (RAC; 0.3 mg/kg; Tocris) was dissolved in 0.9% (w/v) NaCl (saline). GBR12783 dihydrochloride (GBR; 10-20 mg/kg; AdooQ) was dissolved in water. Chloro-APB hydrobromide (1-2 mg/kg, Sigma-Aldrich) was dissolved in saline containing 1% ethanol. All drugs were administered intraperitoneally (i.p.). Prior to testing, mice were habituated to intraperitoneal injections for 3 days. For experiments examining the effects of these drugs on intracellular activation, mice received two injections separated by 15 min. Animals were assigned to one of five treatment groups: Vehicle-Vehicle, RAC-Vehicle, Vehicle-GBR, RAC-GBR, or GBR-RAC. For experiments assessing neural activation by immunofluorescence, mice were perfused 15 min after the second injection, as described below. In some fibre photometry experiments, the same mice received different treatments on separate experimental days in a repeated-measures design.

### Behavioural procedures

#### Apparatus

All training procedures were conducted in sound- and light-attenuating modular operant chambers (Med Associates Inc, PO). Chambers were illuminated by a 3 W, 24 V house light. A recessed food magazine was located in the centre of one wall and connected to a pellet dispenser capable of delivering 20 mg Dustless Precision Reward Pellets (Bio-Serv) into the magazine when activated. Retractable levers were positioned on either side of the magazine. Experimental events, including lever extension and retraction, house light illumination and pellet delivery, were controlled using Med-PC software (Med Associates). The software recorded timestamp events with a temporal resolution of 10 ms, including lever presses, magazine entries and pellet deliveries throughout each session. Behaviour was also monitored using either D-Link IP LED infrared cameras (VGA 1/5 inch CMOS sensor with built-in microphone) or Luxonis PoE cameras with custom recording software.

#### Magazine training

To maintain motivation for food rewards, mice were mildly food deprived and maintained at approximately 85% of their free-feeding weight throughout the experiment. Prior to instrumental training, mice underwent at least 3 days of magazine training in the operant chambers (1 session/day). Each mouse was assigned to a single operant chamber, which was used throughout the experiment. Sessions began with illumination of the house light and ended when the house light was extinguished. Throughout magazine training, both levers remained retracted, allowing mice to freely explore the chamber. Depending on the experiment, 30 or 40 grain-based food pellets were delivered into the magazine according to a random time schedule (RT 60 s) over approximately 30 min. Pellet consumption was verified at the end of each session. For experiments requiring tethering to patch cords (fibre photometry, optogenetics), habituation to the tethering was introduced gradually during the magazine training. On the first day, mice were placed in the chamber without a patch cord attached. On the second day, a single patch cord was connected. Where two patch cords were required, the second patch cord was introduced on the third day, followed by an additional day of magazine training with both patch cords attached.

#### Single-lever instrumental training

These procedures were based on single-lever operant conditioning, in which an action (lever press) was associated with the delivery of an outcome (food pellet). Mice were trained once daily. Each training session began with illumination of the chamber house light and extension of the lever associated with pellet delivery. Sessions ended when the house light turned off and the lever retracted. Assignment of right or left lever was counterbalanced across mice.

#### Random Ratio training

To examine how instrumental learning affected striatal activation, groups of mice (n = 8) underwent a graded training sequence designed to establish and progressively strengthen an action-outcome contingency. Each training session continued until mice had earned 40 pellets or until 60 min had elapsed, whichever occurred first. Following magazine training, mice received four days of continuous reinforcement (CRF), in which every lever press was rewarded. Mice then progressed through a series of random ratio schedules in which the probability of reward (P) delivery following a lever press was gradually reduced across sessions: RR5 (P = 0.2) for 4 days, RR10 (P = 0.1) for 4 days and RR20 (P = 0.05) for 4 or 8 days. Different groups were perfused at specific stages of training to assess changes in striatal activation across learning. One group (first contingency - 1stC) was perfused early during the first day of CRF, 20 min after reaching the first contingency criterion. This criterion required at least two lever presses separated by any interval, followed by two consecutive lever presses occurring less than 60 s apart. Reaching this criterion was defined as the first contingency point, from which a timer was initiated to determine the post-contingency interval before perfusion. A non-contingent control group (NC) was included to control for neuronal activation associated with exposure to the chamber, lever and reward. Following magazine training, these mice received a session in which grain-based food pellets were delivered at random time schedule (RT 60 s), while a dummy lever, counterbalanced for side and without programmed reward delivery, was inserted at the start of the session. The remaining instrumental groups were perfused following completion of CRF, RR5, RR10, RR20, or the extended RR20 schedule (aRR20; 8 days). Finally, a yoked control group (Yoked) was trained for the same duration as the aRR20 group. Each yoked mouse received a pellet whenever its paired “master” mouse earned a reward through lever pressing. Thus, yoked mice experienced the same chamber exposure and reward consumption as the instrumental animals, but reward delivery was independent of their actions and therefore did not support the formation of an action-outcome contingency.

#### Random Interval training

A short random interval (RI) protocol was used in fibre photometry and optogenetic experiments. This schedule of reinforcement was chosen because reinforcement is weakly localised, introducing enduring uncertainty regarding the causal efficacy of any individual response^48,49,69,70^. Following magazine training, mice received one day of CRF, one day of RI15 and three days of RI30 training. Training was conducted on a single-lever, single-outcome basis, with lever identity counterbalanced across animals. Sessions started with illumination of the house light and extension of the lever, and ended after 30 pellet deliveries or 60 min, whichever occurred first. Following the final RI30 session, mice underwent a single-lever outcome-devaluation test. For the biPAC optogenetics experiment, training was extended to establish a second action-outcome contingency using the alternative lever and outcome (purified pellets). Mice received one day of RI15 training followed by three days of RI30 training on the second contingency. This was followed by a two-lever outcome-specific devaluation test.

#### Single lever outcome devaluation test

This procedure followed training on a single action-outcome contingency (original contingency). During the 3 d preceding completion of training, mice were habituated to the devaluation chambers for 30 min per day to minimise novelty-induced stress during testing. Habituation was conducted in a separate room in long, narrow boxes containing an empty plastic food dish positioned on the chamber floor. On test day, mice were placed individually into the devaluation chambers and given 1 h of unrestricted access to either the outcome earned during training (grain pellets) or standard laboratory chow. The order of outcome and chow satiation was counterbalanced across groups. Immediately afterwards, mice were placed in their assigned operant chambers for a 10 min extinction test. The house light was illuminated and the trained lever was available throughout the session, but no outcomes were delivered irrespective of lever pressing. Following a single RI30 retraining session, a second devaluation test was conducted under identical conditions, with satiation conditions reversed.

#### Two-lever outcome-specific devaluation test

This procedure followed training on a second action-outcome contingency (additional contingency). Mice underwent two outcome-specific devaluation tests on consecutive days. Before each test, mice were given 1 h of unrestricted access in the devaluation chambers to either grain pellets or purified pellets, with the order of outcome satiation counterbalanced across groups. Immediately afterwards, mice were placed in their assigned operant chambers for a 10 min extinction choice test. Both levers were available throughout the session, allowing simultaneous assessment of responding on the original (A1-O1) and additional (A2-O2) contingencies in the absence of outcome delivery. On the following day, the alternative outcome was devalued and the procedure repeated.

### Recording of locomotor activity

#### Recording, analysis and visualisation of locomotor activity data

Ambulatory activity was assessed in a rectangular open-field arena (48 x 37.5 cm floor area) with an HD-camera recording from above (Logitech). Mice were habituated to the open-field for 3 consecutive days before testing. Video was collected from the last day of habituation and all test days (1080p, 30-60fps). To extract ambulatory activity from video recordings two approaches were used. For simple tracking of animal position and velocity, the ezTrack video analysis pipeline was used^64^. To visualise this activity, the position coordinates (*x, y*) were extracted and presented as tornado plots. In these plots, the *x*- and *y*-coordinates define spatial location, while the *z*-axis represents time, allowing changes in movement patterns to be visualised throughout the recording session. Alternatively, velocity was plotted in alignment with corresponding neural signals. For more detailed analysis of angular speed and head velocity, videos were analysed using DeepLabCut^65^, with points extracted for both ears (other points obtained but not used). The two ear locations were used to generate a head location and head direction, which were then differentiated to obtain the corresponding speed and angular speed. For synchronisation, LED flashes were triggered by the Med-PC system and time-stamped into the fibre photometry data stream.

### Chemogenetics

To manipulate molecular signalling in D2-SPNs, A2a-Cre::D2-eGFP mice received unilateral stereotaxic injections of a Cre-dependent hM4Di DREADD virus (AAV5-hSyn-DIO-hM4D(Gi)-mCherry). The contralateral hemisphere served as a within-subject control and received an equivalent volume of saline. Three weeks after surgery, mice were habituated to intraperitoneal (i.p.) injections for 3 consecutive days. On the test day, mice received clozapine-N-oxide (CNO; 3 mg/kg, i.p.), followed 30 min later raclopride (0.3 mg/kg, i.p.). Mice were perfused 15 min after the second injection, and brains were processed for immunofluorescence as described below.

### Fibre photometry

To measure neural signals, mice received unilateral injections of viral vectors encoding fluorescent biosensors and underwent optical-fibre implantation. Dopamine was measured in C57BL/6J mice using dLight1.1 (AAV5-CAG-dLight1.1). The intracellular sensors were specifically targeted to D1- or D2-SPNs through Cre-dependent expression in D1-Cre or A2a-Cre mice, respectively. Calcium activity was measured in both D1- and D2-SPNs using GCaMP6m (AAV1.Syn.Flex.GCaMP6m). cAMP levels were measured in D1-SPNs using cADDis (AAV1-hSyn-DIO-cADDis-DW-green). Protein kinase A (PKA) activity was measured in D1-SPNs using ExRai-AKAR2 (AAV2/1-FLEX-syn1-ExRaiAKAR2), with Cre-independent mCherry (AAV5-hSyn-mCherry) co-expressed as an external reference fluorophore.

#### Data Acquisition

Fibre photometry recordings were acquired using Tucker-Davis Technologies (TDT) hardware controlled by Synapse software (TDT). Recordings were performed using either an RZ10x Lux Integrated Processor equipped with integrated LED light sources and a photodetector, or an RZ5P processor coupled to a Doric Lenses LED driver and Newport visible femtowatt photoreceivers. Excitation and emission light were routed through a fluorescence mini cube (Doric Lenses) and delivered to the implanted cannulae via 400 μm core, low-autofluorescence optical patch cords (Doric Lenses). Excitation LEDs were sinusoidally modulated at distinct frequencies: 405 nm at 210 Hz, 465 nm at 330 Hz, and 560 nm at 530 Hz. Fluorescence signals were recovered by real-time lock-in demodulation in Synapse, low-pass filtered at 6 Hz and acquired at a sampling rate of 1017 Hz. For recordings of GCaMP6m, dLight1.1 and ExRai-AKAR2, fluorescence was excited using 405- and 465 nm LEDs, with 20–50 μW delivered at the patchcord face for each wavelength. For cADDis recordings, excitation parameters were modified because the available 405- and 465-nm excitation wavelengths were suboptimal for mNeonGreen. The 465-nm excitation power was therefore increased to approximately 100 μW. To avoid the potential photobleaching and phototoxicity associated with also increasing 405-nm excitation power, mCherry was co-expressed as a reference fluorophore and excited at 560 nm (10-20 μW).

### biPAC optogenetics

#### Optimisation

Initial stimulation parameters were guided by previous studies using biPAC in neuronal populations outside the dorsal striatum^71,72^. Light sensitivity and downstream cAMP signalling could differ in D2-SPNs, so optimal stimulation parameters were determined empirically in A2a-Cre mice expressing Cre-dependent biPAC (AAV5-DIO-EF1a-mKate2-biPAC) with simultaneous fibre photometry recording. Direct measurement of light-evoked cAMP responses was not feasible because cADDis and biPAC showed poor co-expression in D2-SPNs, consistent with substantial viral exclusion. PKA activity was therefore monitored downstream using Cre-dependent ExRai-AKAR2. To improve co-expression, biPAC was injected first, followed one week later by ExRai-AKAR2 injection and optical-fibre implantation.

#### Calibration of optogenetic stimulation with fibre photometry

Combined optogenetic and fibre photometry experiments were performed using an RZ5P processor controlled with Synapse software. Stimulation duration and intensity were configured in Synapse and initiated either manually or by TTL signals generated by the Med-PC system. Two 465 nm LEDs were coupled through a branching patch cord to a single excitation port of the fluorescence mini cube. One LED was operated at low power for photometry recording, whereas the second was operated at higher power to activate biPAC. A 405 nm excitation channel was acquired concurrently as the reference signal, and a single low-autofluorescence patch cord connected the optical system to the implanted fibre while the animal was recorded in an open arena. The continuous 465 nm excitation required for ExRai-AKAR2 photometry was expected to produce a small degree of basal biPAC activation, so excitation power was maintained at the lowest level that provided an adequate fluorescence signal and was held constant throughout each recording (<15 μW at the patch-cord end face). Optogenetic stimulation power was also measured at the patch-cord end face. For intensity-response experiments, 1 s stimuli were delivered at 0.2 mW increments from 0.2 to 1.2 mW, with three repetitions at each intensity. Stimuli were separated by at least 30 s, with intervals extended to as long as 2 min at higher intensities to allow the fluorescence signal to return towards baseline. For duration-response experiments, stimuli of 100 ms, 500 ms, 1 s, and 5 s were delivered three times at each duration. Stimuli were separated by at least 30 s, with intervals extended to 1 min for longer stimulation durations. Responses were averaged across the three repetitions for each condition. Based on these experiments, stimulation at 1 mW for 1 s was selected for subsequent behavioural experiments. To assess responses to a behaviourally relevant stimulation pattern, data from D1-SPN ExRai-AKAR2 recordings collected during the final two days of RI30 training were used to identify a representative session. The selected animal had the median session duration and number of outcomes across the cohort. Outcome times from this session were then used as a temporal template for biPAC activation while PKA activity was monitored using ExRai-AKAR2 fibre photometry.

#### Histological validation of biPAC-induced intracellular signalling

In a first experiment, A2a-Cre::D2-eGFP mice received unilateral expression of Cre-dependent biPAC and contralateral expression of mCherry together with bilateral optical fibre implants. Both hemispheres received a high-frequency stimulation protocol designed to maximise biPAC activation (1 s on, 10 s off for 30 min; 1 mW). The mCherry-expressing hemisphere served as a within-animal control for non-specific effects of optical stimulation, fibre implantation and tissue heating on nuclear signalling. Animals were perfused immediately following completion of the stimulation protocol and brains were processed for immunofluorescence. In a second experiment, A2a-Cre::D2-eGFP mice received bilateral expression of Cre-dependent biPAC, together with bilateral optical fibre implants. One hemisphere received a high-frequency stimulation protocol (1 s on, 10 s off for 30 min; 1 mW), while the contralateral hemisphere was not stimulated and served as a within-animal control. Mice were habituated for 3 consecutive days to the open-field arena, attachment of a single patch cord, and i.p. injections. On the test day, mice were placed in the open-field arena for 10 min period, after which optical stimulation was initiated and maintained for the remainder of the experiment. Chloro-APB (1.5 mg/kg, i.p.) was administered 15 min after stimulation onset, and animals were perfused 15 min later. Brains were then processed for immunofluorescence as described below.

#### Behaviour-driven biPAC stimulation

For behavioural experiments, A2a-Cre mice received bilateral injections of Cre-dependent biPAC and bilateral optical-fibre implants. Stimulation timing was controlled by TTL signals generated by the Med-PC system, which were input to the RZ5P fibre photometry system and then routed to the LED drivers (Doric Lenses). Light was generated by a 465nm rotary LED (Doric Lenses) and delivered to the animal bilaterally via a splitter branching patch cord (Doric Lenses). Reward-triggered stimulation was delivered as a 1 s pulse at 1 mW, based on the prior intensity- and duration-response characterisation experiments. A 10 s time-out was imposed after each stimulation, during which no additional stimulation was delivered even if another reward was earned. For light-blocked conditions, optical output from the patch cord was prevented by coating the fibre end face with black nail polish and filling the ceramic sleeve with plasticine; absence of light output was confirmed before connection to the animal. This preserved the behavioural context of light delivery without stimulating biPAC activity.

### Tissue processing and immunofluorescence labelling

Transcardial fixation and tissue sectioning. Mice were rapidly anesthetized by exposure to 10 sec of isoflurane gas in an induction chamber (4% in air), followed by a lethal intraperitoneal injection of sodium pentobarbital (0.5 mL of 10% Lethabarb, Virbac). After ensuring deep anaesthesia by testing paw and tail reflexes, mice were perfused transcardially using an air pressure system (constant flow of 15 ml/min) with cold formaldehyde solution (4%, w/v) in phosphate buffer saline (pH 7.4). Brains were dissected and post-fixed overnight in 4% formaldehyde solution at 4°C. Consecutive 30 μm coronal sections spanning the rostro-caudal extent of the striatum were obtained for each animal using a vibratome (Leica Microsystems). Free-floating slices were stored at -20°C in an anti-freeze solution (ethylene glycol, 30% v/v; glycerol, 30% v/v, 0.25 M Tris buffer) until processed for immunofluorescence.

#### Immunofluorescence staining

Free floating brain slices (bregma: ∼0.14 mm) were rinsed 3 times in Tris-buffered saline solution (TBS, 0.5 M) for 10 min on an orbital shaker at room temperature (R.T.) prior to membrane permeabilization treatment with triton X-100 solution (0.2%, v/v in TBS), which was applied for 1 h 30 min at R.T. After 3 10-min washes with TBS, slices were incubated at 4°C on an orbital shaker with rabbit polyclonal anti-phospho (Ser10)-Histone H3 (p-H3; 1:400 dilution; Sigma Aldrich); rabbit polyclonal anti-phospho (Ser10)-acetyl (Lys14)-Histone H3 antibody (1:400 dilution; Sigma Aldrich); rabbit polyclonal anti-phospho-ERK1/2 (p-ERK1/2, 1:300 dilution, Cell Signaling Technology) and mouse monoclonal anti-dopamine and cyclic adenosine monophosphate-regulated phosphoprotein, Mr 32 kDa antibodies (DARPP-32; 1:500 dilution; BD Biosciences). Following a 48-h incubation period, unbound primary antibodies were washed off with 4 washes of TBS, and bound primary antibodies were detected through incubation for 1 h 30 min at room temperature with donkey anti-rabbit Alexa Fluor 546 (1:400 dilution; Thermo Fisher Scientific); donkey antirabbit Alexa Fluor 647 (1:400 dilution; Thermo Fisher Scientific); donkey anti-mouse Alexa Fluor Plus 647 (1:400 dilution; Thermo Fisher Scientific) secondary antibodies. Unbound antibodies were washed off through 2 10-min rinses in TBS and 2 10-min final washes in TB. Brain samples were then placed on glass slides and mounted with a coverslip on Vectashield antifade mounting medium (Vector Laboratories, CA). Slides were stored at 4°C in the dark until image acquisition.

### Image acquisition

#### Spinning disk confocal microscopy

Brain sections were imaged using spinning-disk confocal microscopy on Nikon Eclipse Ti-E motorized inverted microscopes equipped with either an Andor Diskovery or a CrestOptics X-Light V3 imaging system and a pco.edge 4.2 sCMOS camera. Automated image acquisition was controlled through Nikon NIS-Elements software, which coordinated microscope operation and motorized stage movement. High-resolution mosaic images spanning each brain hemisphere were generated by automated acquisition of adjacent image tiles within userdefined regions, followed by image stitching to produce a single composite image for analysis.

### Quantification and statistical analysis Instrumental conditioning analysis and visualisation

#### Analysis and visualisation of instrumental conditioning data

Time-stamped datasets were imported into MATLAB (MathWorks, Natick, MA, USA) and Python for analysis and visualisation. To visualise the temporal organisation of responding and the development of action sequences across training, two complementary visualisation approaches were used. Return maps of inter-press intervals (IPIs) were generated for each session by plotting the duration of an interval (IPI_n_) against the duration of the subsequent interval (IPI_n+1_). Data were pooled across animals within each experimental group. For density maps, IPI pairs were binned in logarithmic space and smoothed using a Gaussian kernel to generate two-dimensional probability distributions, with colour intensity reflecting the relative density of observations. In addition, active lever presses were binned across successive training sessions and displayed as colour-density matrices, in which rows corresponded to individual animals and columns to consecutive time bins across training. Subjects were grouped by experimental condition and ordered according to hierarchical clustering based on the similarity of their response patterns, facilitating visual comparison of training-related changes in responding. To visualise response vigour and the temporal organisation of instrumental responding during the devaluation test, raster plots of lever presses on the trained action were generated for each session. Press events were plotted as a function of time and grouped according to experimental group and devaluation condition (valued or devalued). To characterise changes in response rate across the session, lever presses were pooled within each group and condition, binned over time, converted to presses per second, and smoothed to produce continuous response-rate functions.

### Fibre photometry signal processing and analysis

#### Signal processing in pharmacology experiments

With the exceptions described below, all stages of fibre-photometry signal processing were performed using custom code tailored to the photophysical and response properties of each sensor. Because most experiments examined slow signalling dynamics, the relationship between the reference and sensor channels was estimated during a defined pre-injection baseline and extrapolated across the remainder of the recording, rather than estimated across the entire recording as in conventional processing pipelines. This reduced the risk of incorporating drug-evoked responses into the correction model.

For dLight1.1 and GCaMP6m, demodulated 405- and 465-nm fluorescence signals were downsampled to approximately 100 Hz. Because the channels exhibited distinct photobleaching profiles, double-exponential decay curves were fitted to and subtracted from each channel independently before reference-channel fitting. The rationale and validation for this channel-specific photobleaching correction are presented using dLight1.1 recordings in Figure S6A–C. Most experiments included a 30-min pre-injection baseline, which was used to fit the photobleaching curves that were extrapolated across the pharmacological period. For earlier experiments with 10-min baselines, test-day fits were initialised using parameters obtained from the corresponding habituation day recording. Fits were visually inspected and manually refined when necessary to capture the baseline photobleaching trajectory. Photobleaching correction was omitted when it did not improve alignment between the slow components of the two channels. The baseline-fitting and extrapolation procedures are validated in Figure S6D–F. Following photobleaching correction, the 405-nm reference signal was fitted to the 465-nm signal during the pre-injection baseline using first-order least-squares regression. The resulting slope and intercept were applied across the full recording to generate a fitted reference signal. *ΔF/F* was calculated as the difference between the measured 465-nm signal and the fitted reference signal, divided by the fitted reference signal. The resulting trace was z-scored using the mean and standard deviation of the pre-injection baseline.

For ExRai-AKAR2, demodulated 405- and 465-nm fluorescence signals were downsampled to approximately 100 Hz. Separate photobleaching correction was not applied because it did not improve the correspondence between the two channels. The 405-nm signal was fitted to the 465-nm signal over the pre-injection baseline using first-order least-squares regression, and the resulting fit was extrapolated across the full recording. Sensor activity was calculated as the ratio of the measured 465-nm signal to the fitted 405-nm reference signal and expressed as *ΔR/R* relative to the mean baseline ratio (Figure S6K). Because baseline variance was sensitive to sparse spontaneous transients, baseline z-scoring was not used for comparisons across animals. Instead, *ΔR/R* traces were smoothed using a 30-s moving average and normalised to the peak raclopride-evoked response measured in the same animal. This normalisation procedure is explained and validated in Figure S6L–O.

For cADDis, the 465-nm cADDis and 560-nm mCherry fluorescence signals were downsampled to approximately 15 Hz. mCherry was used as an external reference fluorophore to estimate nonspecific fluctuations in the cADDis signal; validation of the correspondence between the slow components of the two channels is presented in Fig S6g,h. Differences in the bleaching profiles of the green and red fluorophores were not adequately accommodated by sequential photobleaching correction and linear reference-channel regression, so the two channels were processed using a simultaneous nonlinear model using the preprocessing component of the behapy analysis pipeline^73^. This model jointly estimated baseline drift and photobleaching in both channels and the component of the cADDis signal associated with fluctuations in the mCherry reference. The rationale and validation for this approach are presented in Figure S6I, J. *ΔF/F* was calculated as the difference between the measured cADDis signal and the fitted reference signal, divided by the fitted reference signal. The resulting trace was z-scored using the pre-injection baseline.

#### Analysis in pharmacology experiments

In dLight1.1, ExRai-AKAR2 and cADDis recordings, analyses focused on slow drug-evoked changes in signalling; therefore a 30-or 60 s moving average was applied before quantification. dLight1.1 responses typically reached a sustained plateau and were quantified as the mean normalised fluorescence from 5–15 min after each injection. ExRai-AKAR2 and cADDis responses varied over the post-injection period; response magnitude and duration were therefore quantified over the specified 10- or 15 min intervals using the positive area under the curve for ExRai-AKAR2 and the negative area under the curve for cADDis. Analysis of GCaMP6m recordings instead focused on discrete calcium transients. Transient peaks were detected from the unsmoothed, z-scored *ΔF/F* trace using a minimum prominence of twice the median absolute deviation of the baseline signal and a minimum interpeak interval of 1 s. Events with peak widths greater than 10 s were excluded as artefacts. Transient frequency was calculated in 1-min bins, then averaged from 5-15 min after each injection.

#### Regression analysis of neural signals and locomotor activity

To examine the relationship between D1-SPN calcium activity and movement, locomotor activity derived from DeepLabCut tracking was compared with GCaMP photometry signals processed using the Behapy analysis pipeline^73^. Head velocity and angular speed were interpolated to the GCaMP sampling times. For each animal and analysis period, ordinary least-squares regression was used to determine the variance in the GCaMP signal explained by these movement parameters across a range of temporal offsets. The optimal offset was defined as that producing the maximum coefficient of determination (R^2^). For the pharmacological comparisons, the model producing the maximum (R^2^) during the day 1 baseline period was applied to the periods following the first and second injections, and the variance explained was calculated for each period.

Signal processing for behaviour. To examine PKA dynamics associated with instrumental behaviour, ExRai-AKAR2 signals were recorded in D1-SPNs during training on a random interval schedule. To emphasise event-locked transients, fluorescence data were processed using a customised ratiometric-signal algorithm implemented in the Behapy analysis pipeline^73^. Event-aligned responses were examined across training days, with their temporal profile used to inform subsequent optogenetic simulation experiments.

### Neuronal activation mapping and quantification

Images were analysed using ImageJ/Fiji software^66^. Images were automatically registered to a designated reference image using the *StackRegJ_ plugin*, which uses key fiducial markers within both the reference and target images to achieve spatial alignment. Manual realignment was carried out in ImageJ/Fiji using the translate and rotate functions. The outer boundary of the striatum and predefined subregions (e.g., areas of high and low viral transduction) were manually delineated using the freehand tool. The area of each region was measured (mm^2^), and the x-y coordinates defining each boundary were extracted. To quantify the number of activated SPNs in D2-eGFP mice and retrieve their location across the striatum, images were automatically thresholded based on p-H3 or p-ERK immunolabelling and individual nuclei were automatically detected using the *Analyze Particle* command in Fiji, which finds the edge of an object (i.e. nucleus), outlines its contour using the wand tool, creates an individual region of interest (ROI), and then determines the coordinates (x, y) of its centre point (centroid). To determine whether an activated SPN was D1- or D2-type, the generated ROIs were overlaid on the eGFP image and the mean grey value (m.g.v) of all the pixels within the selection was then measured. Resulting neuronal data, including position (x, y coordinates), area and mean grey value of the eGFP channel for each detected ROI, were exported in a tab-delimited spreadsheet format and imported into MATLAB. Custom-made scripts were used to obtain the density of activated SPNs (i.e., p-H3 or p-ERK counts/mm^2^) for each SPN population, based on an eGFP m.g.v threshold, which was obtained by averaging the eGFP m.g.v of a number of identified D1- and D2-SPNs. To calculate neuronal density within defined subregions, we used the MATLAB *inpolygon* function, which identifies points located within the polygon delineated by the x-y coordinates of the ROI boundary.

#### Visualisation of neuronal activation patterns

The position of activated neurons within the tissue was digitally reconstructed by combining a line plot defined by the edge points of the striatum with a scatterplot based on the neurons’ centroid coordinates. Individual neurons were colour-coded according to their eGFP mean grey value to distinguish D1- and D2-SPNs. For group-level visualisation, neuronal centroid coordinates from reconstructed hemisections were combined across animals and displayed as stacked scatterplots. Local neuronal density was estimated using kernel density estimation, and individual points were pseudocoloured according to the density of neighbouring observations. This approach highlights regional clustering of activated neurons while preserving the spatial distribution of individual cells. To visualise local variations in the relative abundance of activated D1- and D2-SPNs, neuronal centroid coordinates were projected into a common spatial reference frame and divided into a regular spatial grid. For each grid element, local D1- and D2-SPN counts were quantified and expressed as the log_2_-transformed D2/D1 ratio. Ratio maps were generated for each coronal section and subsequently averaged across sections to produce a single spatial representation for each animal and treatment condition. Colour intensity reflects the local predominance of activated D2- or D1-SPNs within neighbouring neuronal populations, with positive values indicating D2 predominance and negative values indicating D1 predominance.

### Statistical analysis

Statistical analyses were performed using IBM SPSS Statistics (IBM) and R (R Foundation for Statistical Computing). Detailed statistical information for all analyses is provided in Table S1, including sample sizes, experimental units, statistical tests, test statistics, degrees of freedom, effect sizes, confidence intervals, multiple-comparison procedures, corrections for violated assumptions, and exact P values where available. All statistical tests were two-sided and an α level of 0.05 was used throughout. Differences between two conditions were assessed using paired or independent-samples t-tests, as appropriate. Comparisons involving a single factor were analysed using one-way analysis of variance (ANOVA). When homogeneity of variance assumptions were violated, as assessed by Levene’s test, Welch’s ANOVA was used and Games-Howell post hoc comparisons were performed. Experiments involving repeated measurements were analysed using repeated-measures ANOVA, whereas experiments incorporating both within-subject and between-subject factors were analysed using mixed ANOVA. For repeated-measures analyses with more than two levels, sphericity was assessed using Mauchly’s test. When sphericity assumptions were violated, Greenhouse-Geisser-corrected degrees of freedom and associated P values are reported. Where appropriate, post hoc comparisons were conducted using Bonferroni-corrected tests, Games-Howell tests, least significant difference (LSD) tests, or statistically planned simple-effects analyses, as specified in Table S1. Exact P values are reported wherever possible; values below reporting limits are reported as P < 0.001. Effect sizes are reported as partial η^2^ for ANOVA-based analyses and Cohen’s d for t-tests. Sample sizes (*n*) and experimental units are reported in Table S1 and the corresponding figure legends. Depending on the analysis, the experimental unit comprised biologically independent animals, hemispheres, or tissue sections, as indicated.

## Supporting information

Table S1 - statistics

## Acknowledgements

We thank S Zhang, M Andermann and J Zhang for providing sequence information for the biPAC and ExRai-AKAR2 constructs. We thank R Lawler and R Barnes for assistance with animal care. We thank S Killcross and B Balleine for support of this research, and all members of the Decision Neuroscience Laboratory for discussions.

