## Supplemental information for "Striatal D2-neurons gate D1-mediated associative encoding under uncertainty"

### Supplemental figures

#### Figure S1

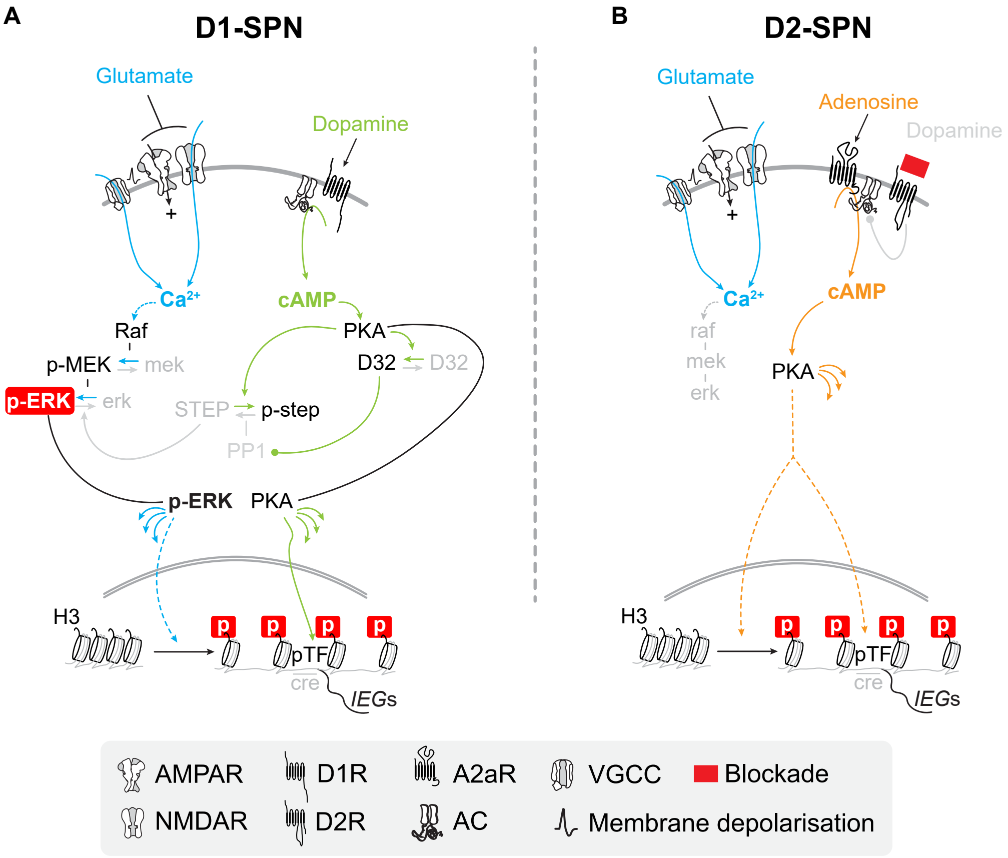


**Figure S1. Distinct intracellular signalling cascades in D1- and D2-SPNs.** Calcium ($Ca^{2+}$) and cyclic AMP (cAMP) initiate broadly similar signalling programs in D1- and D2-SPNs. However, while glutamatergic transmission strongly influences $Ca^{2+}$ activity in both neuronal types (blue), protein kinase activities and nucleosomal responses are engaged through distinct pathways and opposing dopaminergic signals[37,46,54,74].

(A) In D1-SPNs, dopamine activates cAMP signalling through D1-receptors (green), enabling propagation of PKA signalling and amplification of the $Ca^{2+}$-related MAP kinase cascade (p-ERK) via DARPP-32 modulation (D32). Both PKA and p-ERK contribute nucleosomal activation in the nucleus (p-H3).

(B) In contrast, in D2-SPNs, the absence of dopamine lifts D2R-mediated inhibition, allowing adenosine-bound A2a receptor (A2aR) signalling to drive cAMP production and PKA signalling to the nucleus (orange). This pathway does not activate p-ERK in D2-SPNs but can robustly induce nucleosomal responses (p-H3). Thus, p-ERK serves as a reporter of dopaminergic influence in D1-SPNs, whereas pH3 reflects opposing dopamine modulations in both D1- and D2-SPNs.

p: phosphorylated; pTF: phosphorylated transcription factor; IEGs: immediate-early genes.

### Figure S2

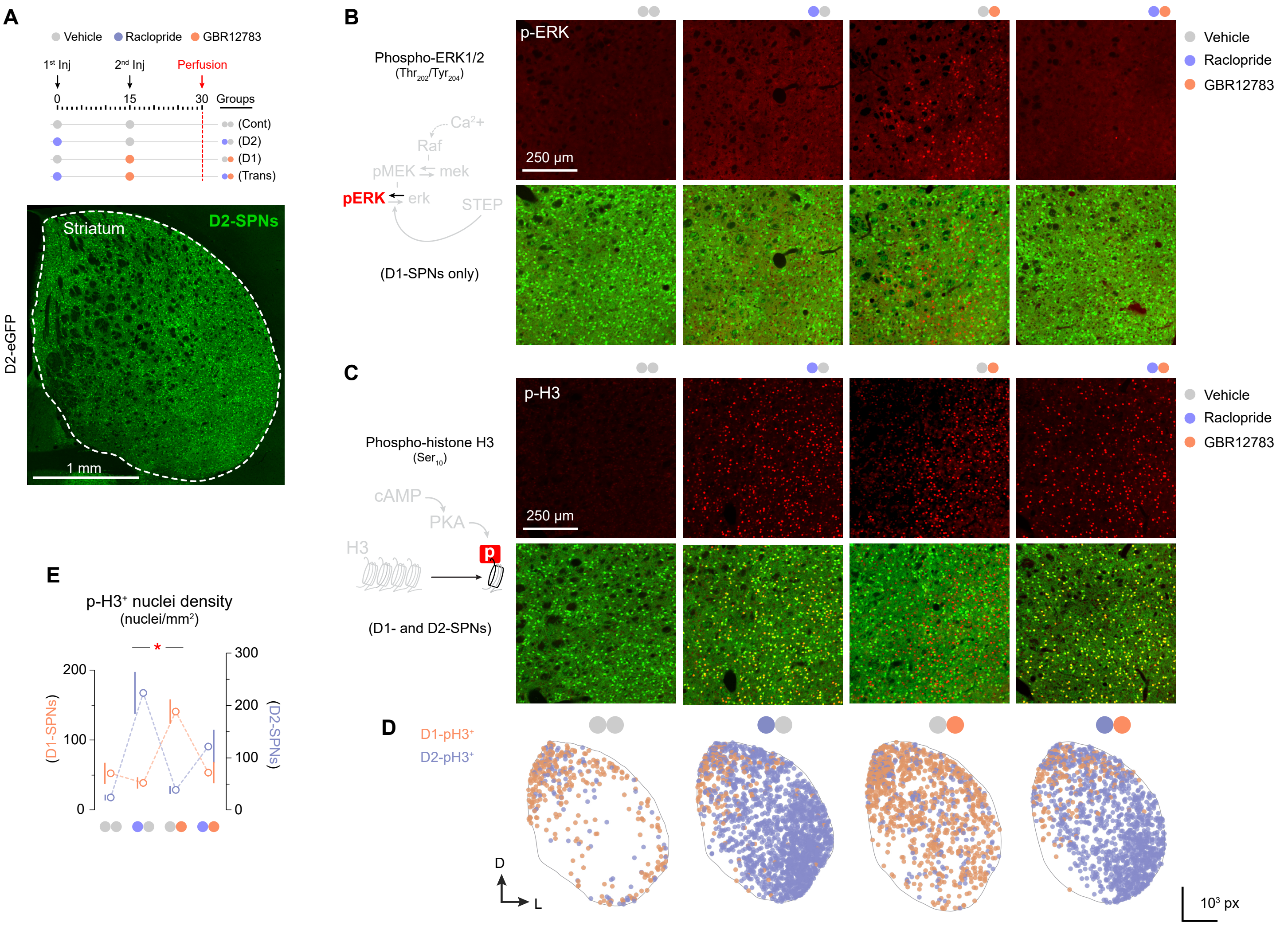


**Figure S2. Striatum-wide D2→D1-SPN transmodulation induced pharmacologically (relates to Figure 2).**
(A) Pharmacological protocol used to induce D2→D1-SPN transmodulation in D2-eGFP mice (right; genetically labelled D2-SPNs). Mice received an initial i.p. injection (1st inj; vehicle or the D2-receptor blocker raclopride) followed by a second injection (2nd inj; vehicle or the dopamine transporter blocker GBR12783). Groups "D2" and "D1" selectively activated D2- and D1-SPNs, respectively. In group "Trans", both striatal systems were activated sequentially, thereby modelling D2→D1-SPN transmodulation.
(B, C) Wide-field confocal micrographs showing (A) phospho-ERK1/2 ($Thr_{202}/Tyr_{204}$) and (C) phospho-histone H3 ($Ser_{10}$) immunofluorescence in the striatum of D2-eGFP mice treated as in (A) and perfused 15 min after the second injection (referred to as p-ERK and p-H3, respectively).
(D) Scatter plots showing spatial distributions of p-H3-positive neurons in D1-SPNs (orange) and D2-SPNs (purple) under each treatment condition (representative hemisections).
(E) Density of p-H3-immunopositive neurons phenotyped as D1-SPNs (left axis) or D2-SPNs (right axis) based on the absence or presence of D2-eGFP fluorescence. Note that in the Trans group, prior administration of raclopride reduced the effects of GBR12783 on D1-SPNs.
See Figure S1 for details of intracellular signalling processes in both SPN subtypes. *, significant interaction (red). See Table S1 for sample sizes and statistical analyses.

### Figure S3

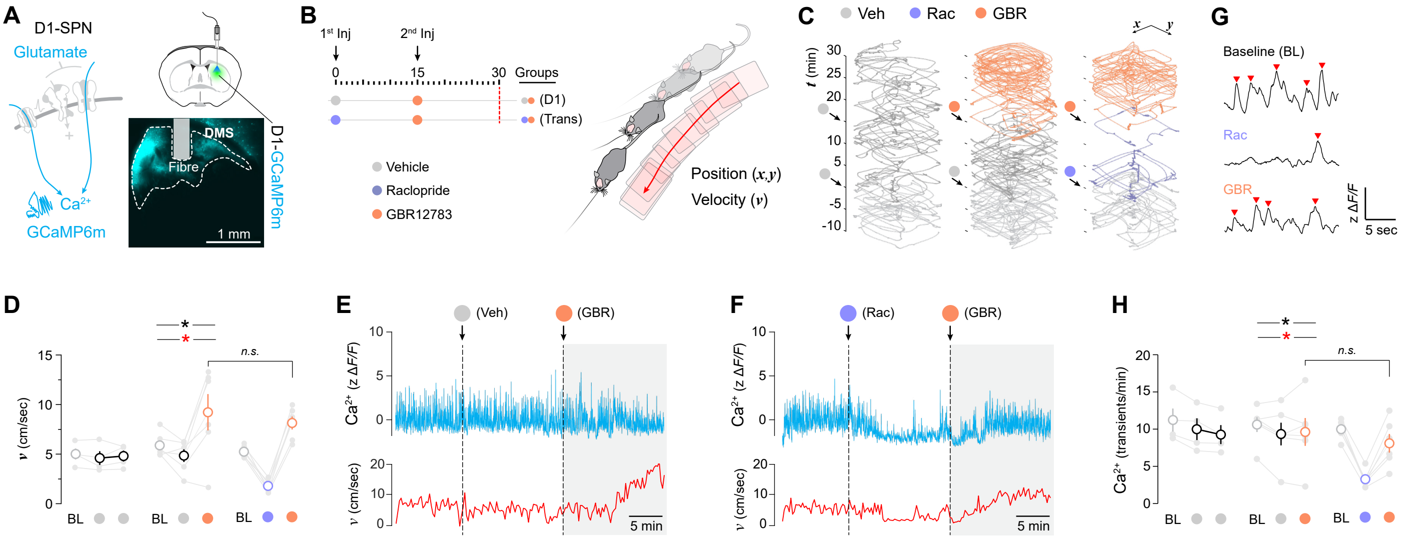

**Figure S3. D1-SPN $Ca^{2+}$ dynamics are insensitive to transmodulation and closely track locomotion (relates to Figure 2).**

(A) Recording of $Ca^{2+}$ dynamics in DMS D1-SPNs (Flex-GCaMP6m) of D1-Cre mice. Right, example injection site with marked fibre tract.

(B) Pharmacological protocol used to induce D2→D1-SPN transmodulation. The injection sequence defined three locomotor activity epochs across the session: baseline, post-first injection, and post-second injection. Right, locomotor activity was quantified in an open field by extracting velocity derived from *x,y* video tracking of mouse position.

(C, D) Example tracked trajectories (C) and quantified velocity (D) of mice receiving different pharmacological treatments.

(E, F) Examples of D1-SPN $Ca^{2+}$ activity (blue) and corresponding locomotor velocity (red) in animals receiving Veh and GBR (E) or Rac and GBR (F).

(G) Peak detection procedure used to identify $Ca^{2+}$ transients from Flex-GCaMP6m recordings. Red arrowheads indicate detected events exceeding threshold.

(H) Comparison of $Ca^{2+}$ transient rates (transients/min) across treatment groups and experimental epochs.

Data in (D) and (H) show individual values (light grey), mean values (dots) and SEM (error bars). *, significant overall effect (black) and interaction (red). *n.s.*, not significant. See Table S1 for sample sizes and statistical analyses.

### Figure S4

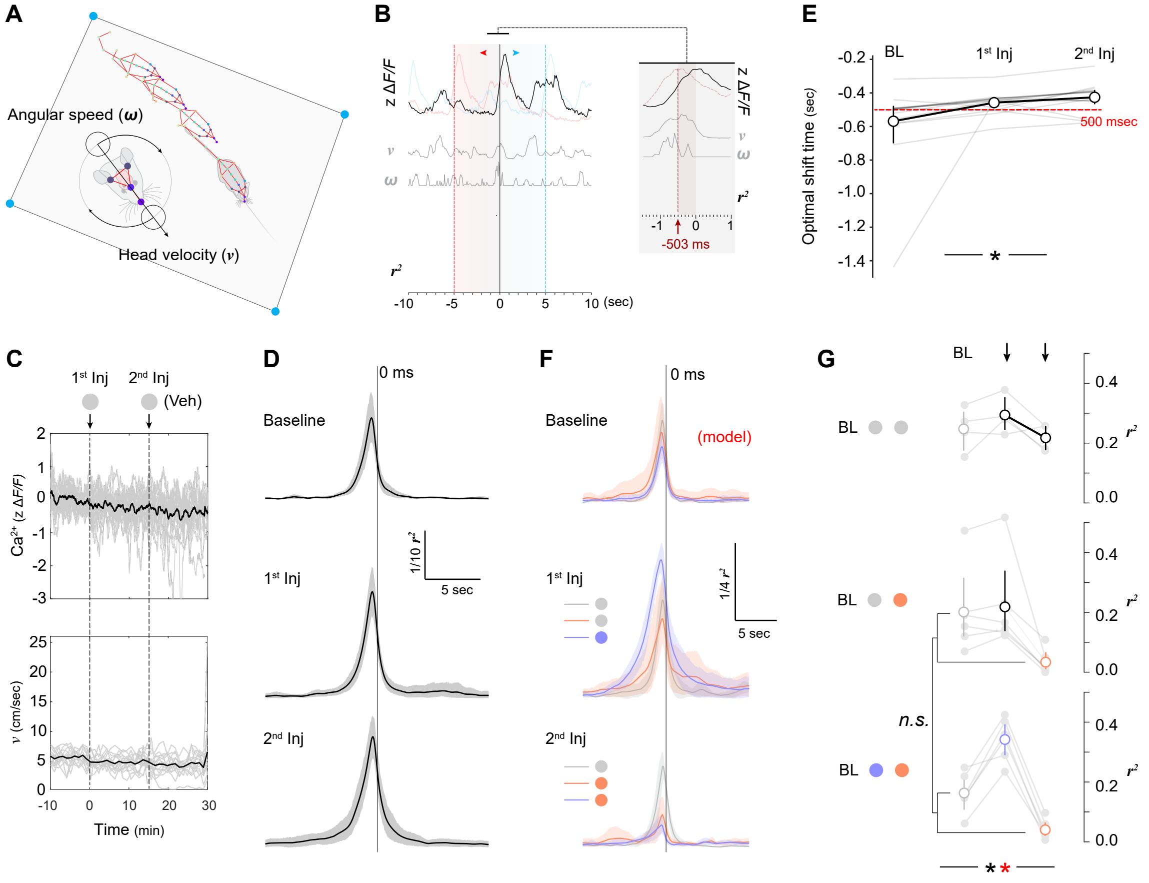


**Figure S4. D1-SPN $Ca^{2+}$ dynamics consistently succeeds behavioural correlates irrespective of pharmacological state (relates to Figure 2).**

(A) Animal pose analysis performed on video recordings of pharmacologically treated mice freely moving in an open field was used to calculate angular speed ($\omega$) and head velocity ($v$).

(B) Head motion parameters ($\omega$ and $v$) were regressed against temporally shifted $Ca^{2+}$ signals simultaneously recorded in DMS D1-SPNs (anterograde shift, blue; retrograde shift, red) to determine the optimal regression time ($r^2$). Inset, expanded view of a representative alignment between head motion and corresponding neural activity peaks (approximately 500 ms).

(C) DMS D1-SPN fibre photometry recordings (top) and corresponding locomotor velocity (bottom) in D1-Cre mice during the third day of habituation to the open-field chamber. Vehicle (saline) injections were administered at the indicated times, defining three analysis epochs (baseline, first injection and second injection).

(D) Optimal regression time between head motion parameters and temporally shifted $Ca^{2+}$ signals across each epoch. Vertical line denotes zero shift. Behaviour consistently preceded corresponding $Ca^{2+}$ signals.

(E) Optimal regression times obtained from individual animals across epochs. All three epochs yielded comparable values of approximately 500 ms.

(F) In animals treated as in Figure 2d (first injection: vehicle or raclopride; second injection: vehicle or GBR12783), the optimal regression time determined during the baseline period ($t = -10$ to 0 min) was applied to subsequent post-injection epochs (first and second injection) in each pharmacological group. Vertical line denotes zero shift.

(G) Comparison of the strength of head motion-$Ca^{2+}$ activity regressions across groups and treatment epochs.

Data in (E) and (G) show individual values (light grey), mean values (open circles) and bootstrapped confidence intervals (error bars). *, significant overall effect (black) and interaction (red). *n.s.*, not significant. See Table S1 for sample sizes and statistical analyses.

#### Figure S5

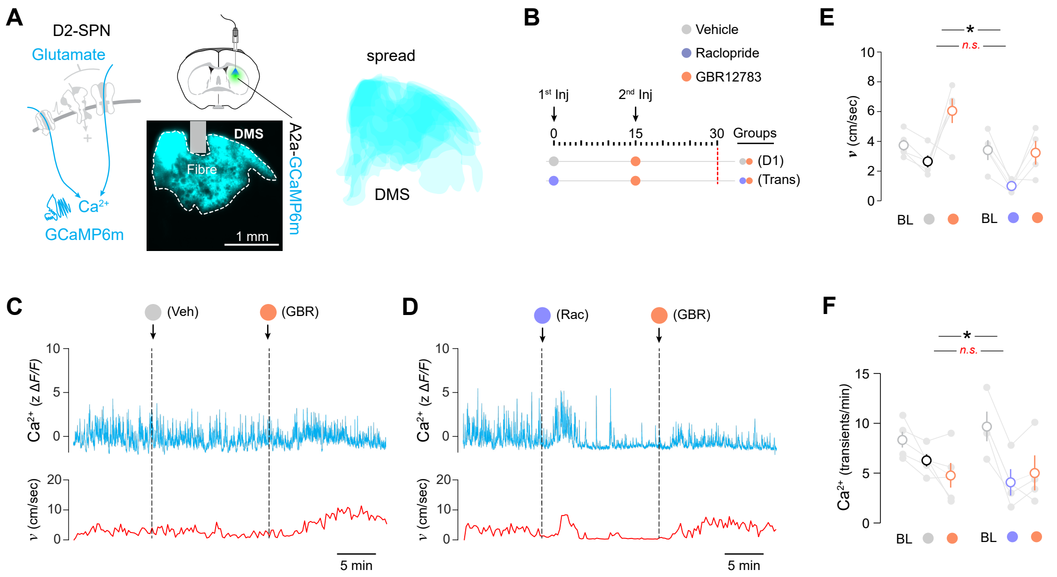


**Figure S5. $Ca^{2+}$ dynamics in D2-SPNs follows locomotion and does not report transmodulation (relates to Figure 2).**
(A) Recording of $Ca^{2+}$ dynamics in DMS D2-SPNs (Flex-GCaMP6m) of A2a-Cre mice. Bottom, example injection site with marked fibre tract. Right, overall DMS expression spread across all mice.
(B) Drug injection schedule used to induce D2→D1-SPN transmodulation. Injections defined three locomotor activity epochs across the session (baseline, first injection and second injection).
(C, D) Examples of D2-SPN $Ca^{2+}$ activity (blue) and corresponding locomotor velocity (red) in animals receiving Veh and GBR (C) or Rac and GBR (D).
(E, F) Comparison of locomotor velocity (E) and $Ca^{2+}$ transients/min (F) across treatment groups and experimental epochs. Data in (E) and (F) show individual values (light grey), mean values (dots) and SEM (error bars). *, significant overall effect. *n.s.*, not significant. See Table S1 for sample sizes and statistical analyses.

### Figure S6

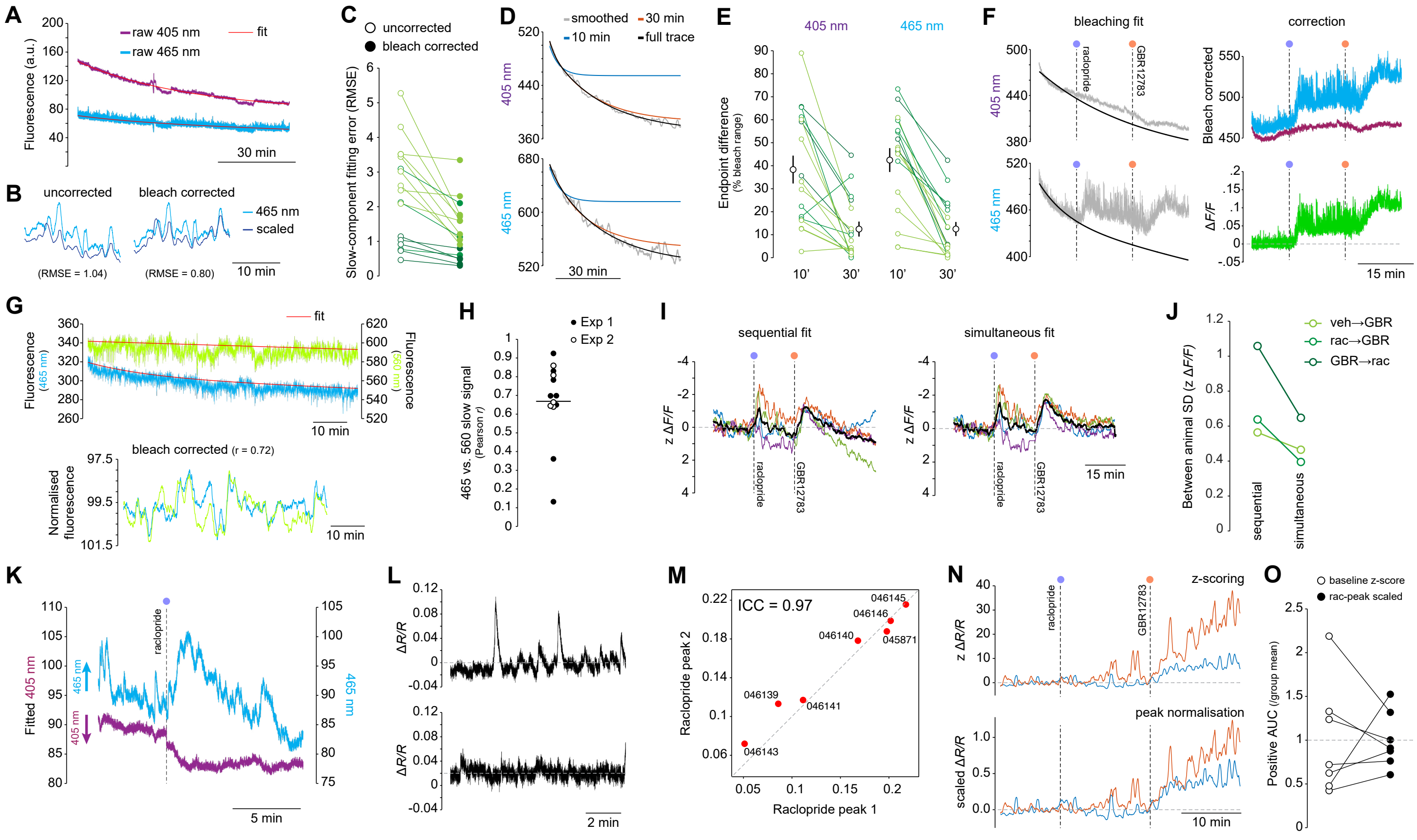


**Figure S6. Analytical approaches used to quantify slow signalling dynamics in fibre photometry recordings (relates to Figure 3).** Standard fibre photometry workflows were adapted to preserve and quantify the slow signalling dynamics associated with pharmacological manipulation and intracellular biosensors. Key processing steps were validated for dLight1.1 (A-F), cADDis (G-J) and ExRai-AKAR2 (K-O).

(A-C) Bleaching correction improves estimation of slow signal components in dLight1.1 recordings. (A) Representative raw 405-nm and 465-nm fluorescence signals recorded during a drug-free habituation session, with independently fitted double-exponential bleaching curves. (B) Expanded example showing linear fitting of the 405-nm reference channel to the 465-nm signal before and after bleaching correction. Correction improved fit quality, as indicated by reduced root mean square error (RMSE). (C) Paired comparison showing reduced slow-component RMSE following bleaching correction across all recording cohorts (n = 17 animals; paired Wilcoxon signed-rank test, W = 153, P < 0.0001).

(D-F) Evaluation of bleaching-curve fitting strategies for dLight1.1 recordings when only limited baseline periods were available. (D) Representative smoothed traces showing double-exponential bleaching curves fitted to either an initial 10- or 30-min baseline period and extrapolated throughout the recording, or fitted to the full recording. (E) Absolute differences between extrapolated and full-recording bleaching fits measured at recording endpoints. Fits based on 10-min baselines were frequently unstable, whereas 30-min baselines showed improved agreement. (F) Example processing workflow for pharmacological recordings with short baseline periods. Bleaching curves estimated from habituation sessions were applied to test-day recordings before calculation of *ΔF/F*. Correction quality was verified manually.

*(legend continued on next page)*

(G, H) Validation of mCherry as an external reference fluorophore for cADDis recordings, given the suboptimal performance of 405-nm excitation as a reference channel. (G) Representative fluorescence traces with independently fitted bleaching curves (top) and corresponding baseline-normalised slow signal components overlaid after correction (bottom). (H) Correlation between normalised 465-nm and 560-nm slow signal components across two independent experiments. Data points represent individual animals; horizontal line indicates the group mean.
(I, J) Simultaneous fitting of signal and control channels reduces analyst intervention and decreases between-animal variability in cADDis recordings. (I) Representative *ΔF/F* traces processed using either sequential bleaching correction followed by linear fitting (left) or a simultaneous model that jointly fits signal and control channels without requiring independent bleaching curves (right). Simultaneous fitting reduced divergence between animals towards the end of recordings and minimised potential fitting bias. Coloured traces represent individual animals; black traces represent group averages. (J) Between-animal variability across pharmacological treatment groups was lower following simultaneous than sequential fitting.
(K-O) Normalisation of ratiometric ExRai-AKAR2 recordings for between-animal comparisons. (K) Representative fitted 405-nm and 465-nm traces used to calculate the ratiometric *ΔR/R* signal. (L) Representative baseline *ΔR/R* recordings from two animals. Sparse stochastic baseline transients produced substantial differences in baseline variance, limiting the suitability of conventional z-scoring approaches. (M) Raclopride-evoked responses were highly reproducible within animals (intraclass correlation coefficient = 0.97) and were therefore used as an internal normalisation reference. (N) GBR12783-evoked responses from the animals shown in (L), normalised using either conventional baseline z-scoring (top) or raclopride-evoked peak responses (bottom). Peak normalisation improved comparability across animals. (O) Positive area-under-the-curve (AUC) measurements of GBR12783-evoked responses expressed relative to the group mean following either baseline z-scoring or raclopride-peak normalisation. Peak normalisation reduced between-animal variability, decreasing the coefficient of variation from 0.633 to 0.320 (CV ratio = 0.505, 95% bootstrap confidence interval = 0.327-0.884; bootstrap P = 0.0187; n = 7).

#### Figure S7

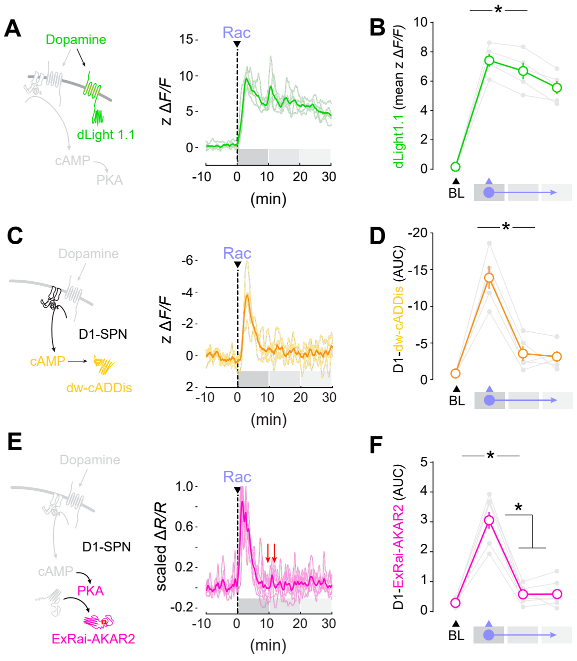


**Figure S7. Raclopride alone triggers D2→D1-SPN transmodulation (relates to Figure 3).** Mice were implanted with optic fibres into the dorsomedial striatum (DMS) for *in vivo* recording of extracellular dopamine (dLight1.1), intracellular cyclic AMP (D1-dw-cADDIs) and protein kinase A activity (D1-ExRai-AKAR2) in D1-SPNs, as in Figure 3.

(A-F) Fibre photometry recordings of dopamine (A, B), cAMP (C, D) and PKA activity (E, F) before and after i.p. administration of the D2-receptor antagonist raclopride (Rac). (A, C, E) Recordings before and after Rac administration. (B, D, F) Quantification of dopamine, cAMP and PKA activity signals across recording periods. Double red arrows in (E) denote an unanticipated drop in ongoing PKA activity, similar to those observed in Figure 3F and L.

Data in (B), (D) and (F) show individual values (light grey), mean values (dots) and SEM (error bars). *, significant overall/simple effect. *n.s.*, not significant. See Table S1 for sample sizes and statistical analyses.

### Figure S8

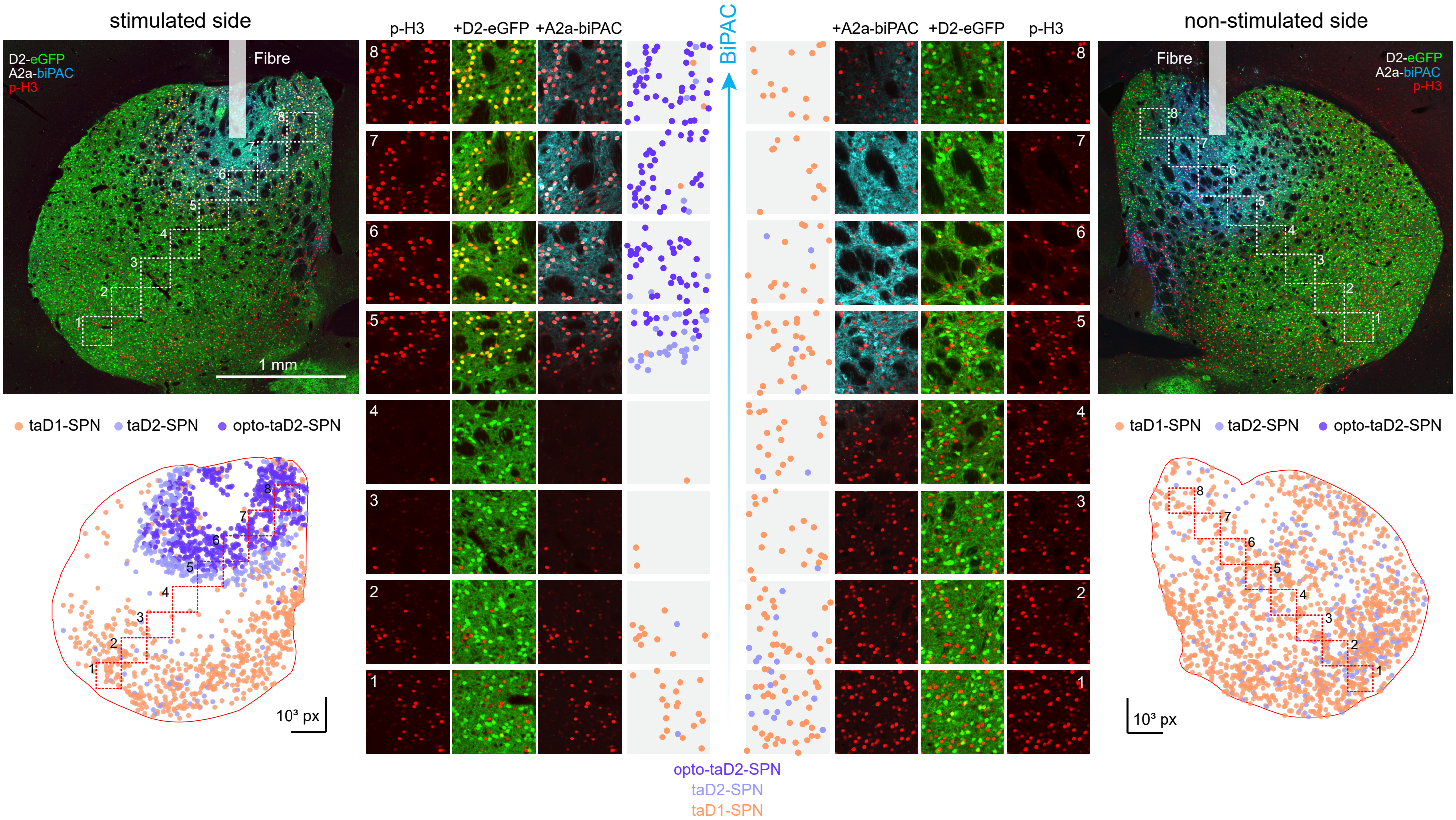


**Figure S8. Optogenetic stimulation of D2-SPN cAMP signalling causes widespread reduction of intracellular signalling in D1-SPNs (relates to Figure 4).** A2a-Cre x D2-eGFP mice bearing bilateral Cre-dependent biPAC expression and optic fibre implants in the DMS received unilateral intermittent optical stimulation and systemic D1-receptor agonist injections before processing for immunofluorescence, as in Figure 4C (stimulation: 1 sec ON, 10 sec OFF, $t$ = 10-40 min; injection: 1.5 mg/kg chloro-APB, $t$ = 25 min). Images show low- and high- magnification (insets 1-8) confocal micrographs of the stimulated (left) and non-stimulated (right) striata from the same animal. Triple labelling identifies transcriptionally active SPNs (taSPNs, p-H3 positive), D1- (eGFP negative) and D2 (eGFP positive)-SPNs, and biPAC-expressing D2-SPNs. Bottom panels show digital reconstructions of taSPNs detected in the corresponding hemisections. Based on fluorescence profiles, taSPNs were classified as taD1-SPNs (no D2-eGFP or A2a-biPAC fluorescence), taD2-SPNs (D2-eGFP fluorescence without observable A2a-biPAC fluorescence) or opto-taD2-SPNs (detectable D2-eGFP and A2a-biPAC fluorescence). A substantial proportion of taD2-SPNs with subthreshold A2a-biPAC signal surrounding the viral spread area are likely to have been optically stimulated. Neurons immediately adjacent to the optic fibre display little intracellular signalling, probably owing to local tissue disruption associated with implantation. Notably, inhibition of D1-SPN signalling extends beyond regions containing detectable A2a-biPAC expression. Central scatterplots show classified data points corresponding to each higher-magnification inset. All images and reconstructions are from the same animal. Quantified effects across animals are shown in Figure 4E-G.

### Figure S9

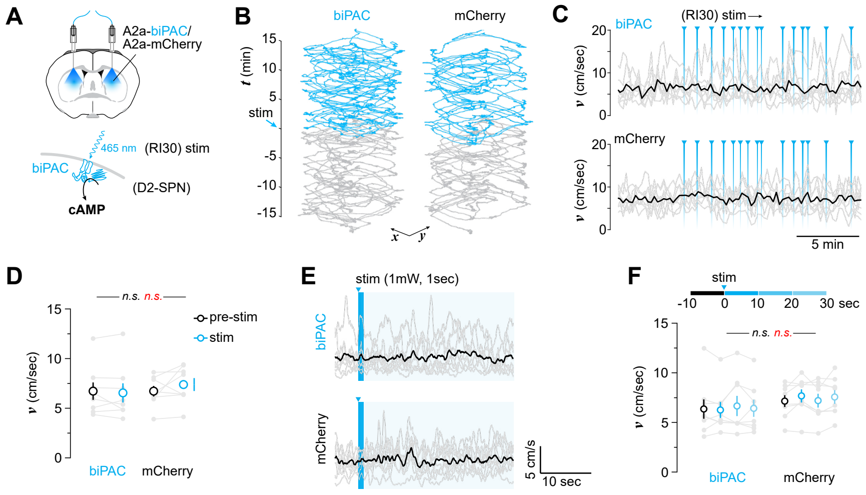


**Figure S9. Optogenetic stimulation of cAMP signalling in DMS D2-SPNs does not affect locomotion (relates to Figure 5).**

(A) Two weeks after completion of the behavioural procedures shown in Figure 5, the same A2a-Cre mice bearing Cre-dependent biPAC or mCherry and bilateral optic fibre implants in the DMS (allowing *in vivo* stimulation of cAMP signalling in D2-SPNs) were exposed to an optogenetic stimulation session in an open field, as in Figure S3.

(B) Examples of locomotor activity before (grey) and during (blue) periods of optogenetic stimulation in a biPAC-expressing and an mCherry control mouse. Traces show tracked *x,y* position over time (*t*).

(C) Individual mice (light grey) and average (black) locomotor velocity (*v*) before and during blue-light stimulation. The same stimulation template was applied to all mice over 30 min and matched the reinforcement frequency and variance previously generated during instrumental RI30 training (Figs. 4 and 5).

(D) Quantification of locomotor velocity during 15 min pre-stimulation and stimulation periods.

(E) Instantaneous velocity (cm/sec) aligned to each stimulation pulse in both groups. Data show averages for each animal across the 15 stimulations (grey) and grand average (black).

(F) Quantification of velocity averaged across 10-sec segments occurring within and outside stimulation periods.

Data in (D) and (F) show individual values (light grey), mean values (dots) and SEM (error bars). *n.s.*, not significant overall effect (black) and interaction (red). See Table S1 for sample sizes and statistical analyses.

**Table S1**
